# Dynamic Interrogation of Stochastic Transcriptome Trajectories Using Disease Associated Genes Reveals Distinct Origins of Neurological and Neuropsychiatric Disorders

**DOI:** 10.1101/2022.02.26.482124

**Authors:** Theodoros Bermperidis, Simon Schafer, Fred H Gage, Terry Sejnowski, Elizabeth B Torres

## Abstract

1

The advent of open access to genomic data offers new opportunities to revisit old clinical debates while approaching them from a different angle. We examine anew the question of whether psychiatric and neurological disorders are different from each other by assessing the pool of genes associated with disorders that are understood as psychiatric or as neurological. We do so in the context of transcriptome data tracked as human embryonic stem cells differentiate and become neurons. Building upon probabilistic layers of increasing complexity, we describe the dynamics and stochastic trajectories of the full transcriptome and the embedded genes associated with psychiatric and/or neurological disorders. From marginal distributions of a gene’s expression across hundreds of cells, to joint interactions taken globally to determine degree of pairwise dependency, to networks derived from probabilistic graphs along maximal spanning trees, we have discovered two fundamentally different classes of genes underlying these disorders and differentiating them. One class of genes boasts higher variability in expression and lower dependencies (“active genes”); the other has lower variability and higher dependencies (“lazy genes”). They give rise to different network architectures and different transitional states. Active genes have large hubs and a fragile topology, whereas lazy genes show more distributed code during the maturation toward neuronal state. Lazy genes boost differentiation between psychiatric and neurological disorders also at the level of tissue across the brain, spinal cord, and glands. These genes, with their low variability and asynchronous ON/OFF states that have been treated as gross data and excluded from traditional analyses, are helping us settle this old argument at more than one level of inquiry.

**Manuscript Contribution to the Field:** There is an ongoing debate on whether psychiatric disorders are fundamentally different from neurological disorders. We examine this question anew in the context of transcriptome data tracked as human embryonic stem cells differentiate and become neurons. Building upon probabilistic layers of increasing complexity, we describe the dynamics and stochastic trajectories of the full transcriptome and the embedded genes associated with psychiatric and/or neurological disorders. Two fundamentally different types of genes emerge: “lazy genes” with low, odd, and asynchronous variability patterns in expression that would have been, under traditional approaches, considered superfluous gross data, and “active genes” likely included under traditional computational techniques. They give rise to different network architectures and different transitional dynamic states. Active genes have large hubs and a fragile topology, whereas lazy genes show more distributed code during the maturation toward neuronal state. Under these new wholistic approach, the methods reveal that the lazy genes play a fundamental role in differentiating psychiatric from neurological disorders across more than one level of analysis. Including these genes in future interrogation of transcriptome data may open new lines of inquiry across brain genomics in general.

## 3 Introduction

The question of whether a distinction should exist between psychiatric and neurological disorders predates the time when psychiatry was not even a formal discipline as we know it today. Back then, motor movements were used as criteria to identify mental disorders, by observing and describing patients in a motor-informed way (*e.g.*, catatonic, hyperactive, *etc.*) [1]. Under the spell of Freud’s psychoanalyses and following Descartes’s dualism, this type of physical-motor criterion lost influence in favor of elaborate descriptions of mental and emotional states inferred by other, non-motor-based criteria. There was more judgment added on to the perception of the patient; for example, terms such as deviant, opposing, defiant, socially inappropriate, behaviors, *etc.,* entered the descriptions of children with atypical neurodevelopment. This judgment took place solely based on external observation, without any additional description of internal states of their nervous systems.

The distinction broadened between mental illness and disorders that affected the person’s function beyond dysfunction of the central nervous system, thus prolonging the ongoing physiological and medical debates [2]; it also impacted the perception of other members of society with regards to one or the other [3]. A recent revival of this debate underscores the side of the argument that psychiatric disorders are not just “mental” but are physical, too, identifying neurobiological substrates of mental illness [4]. These substrates are in line with the current US National Institute of Mental Health Research Domain Criteria (RDoC), a framework that cuts across research domains [5] but still has room for improvement [6; 7; 8; 9; 10].

An example of the brain’s affected tissues that are amenable to separating psychiatric from neurological disorders is provided in Figure 1A and results from this recent resurgence of the debate [11]. Yet, others have contested such neuroimaging distinctions between the disorders, on the grounds that medications can alter brain structures [12]. Specifically, the argument is centered on the ambiguity that medication brings to the studies that are based on brain structure by, for example, increasing basal ganglia volume or increasing volume loss in general, in the case of traditional antipsychotics [12].

**Figure 1.**
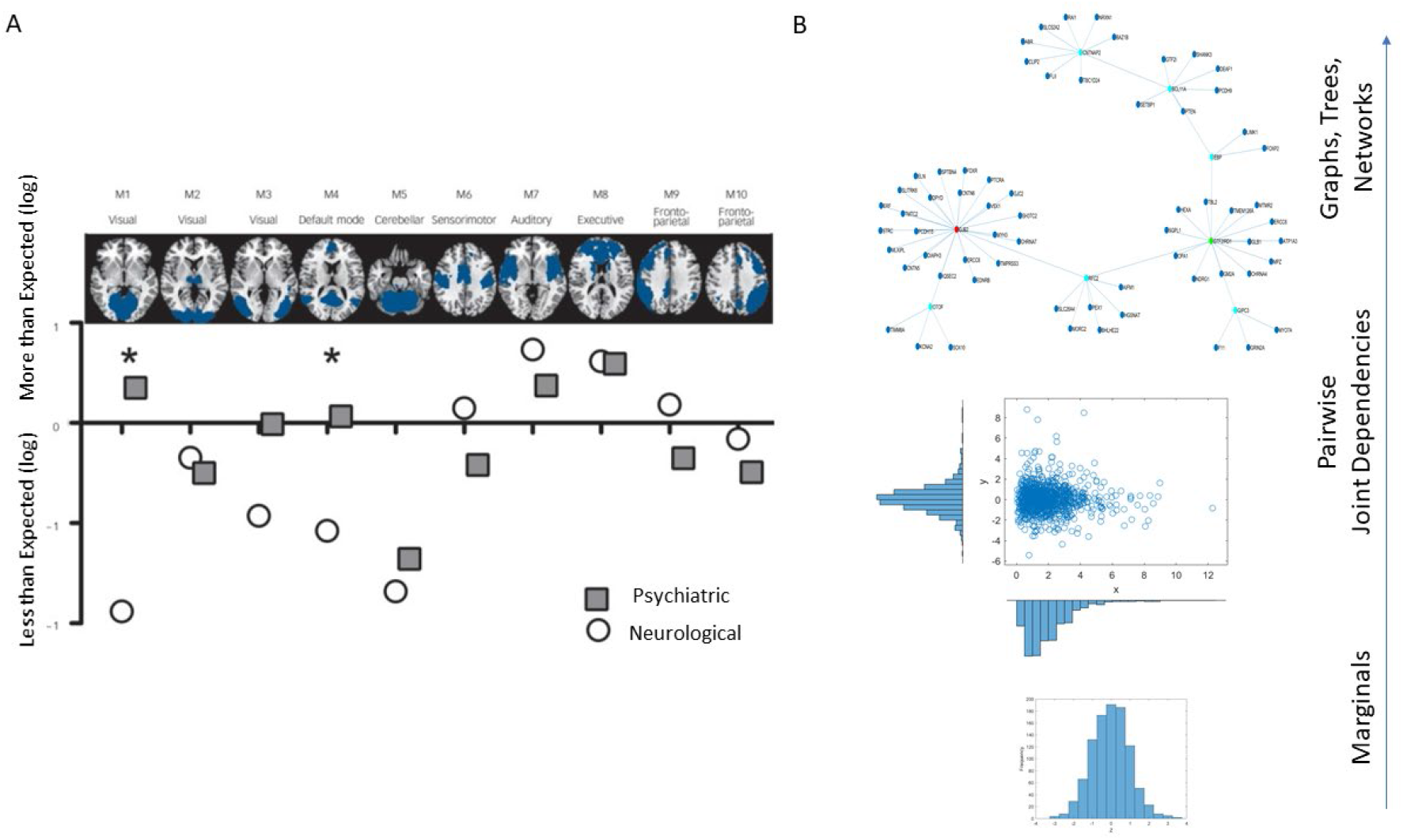
Different lines of inquiry to study the differentiation between psychiatric and neurological disorders. (A) Brain structural differentiation reported by Crossley et al. (Figure from [11] with permission.) (B) Approach used in this work, from simpler to more complex levels of interrogation and the consideration of important dynamically changing statistical co-dependencies across genes’ expressions. Marginal distributions of different genes’ expression across cells followed by studies of pairwise genes’ relations and evaluation of degree of genes’ co-dependendies in joint distributions, followed by analyses of complex, dynamically changing, interconnected networks of genes, across days of embryonic stages of cell differentiation toward neuronal states.

The interactions between diagnosis and medication are also mentioned in the Diagnostic Statistical Manual DSM-5 as a way to justify the exclusion of motor criteria from diagnosis “*Medication-induced movement disorders are included in Section II because of their frequent importance in (1) the management by medication of mental disorders or other medical conditions and (2) the differential diagnosis of mental disorders (e.g., anxiety disorder versus neuroleptic-induced akathisia; malignant catatonia versus neuroleptic malignant syndrome). Although these movement disorders are labeled ‘medication induced’, it is often difficult to establish the causal relationship between medication exposure and the development of the movement disorder, especially because some of these movement disorders also occur in the absence of medication exposure. The conditions and problems listed in this chapter are not mental disorders.*” (Emphasis added.) Nevertheless, several of the mental disorders on a spectrum, like autism spectrum disorder (ASD), attention deficit hyperactivity disorder (ADHD) and schizophrenia do have functional neuromotor issues with neurobiological bases, *i.e.,* of the neurological type, even when medication was never used [13] or was sparsely used [14]. Thus, the confounds between medication and psychiatry- or neurology-based diagnoses are palpable at the clinical level and confusing at the level of basic brain research.

One avenue that we could explore to try and distinguish psychiatric and neurological disorders is by re-examining brain (and bodily tissues) from the standpoint of dynamically changing gene expression in early embryonic stages of pluripotency, as cells transition into neuronal classes. In this context, we could use different levels of inquiry. For example, we could interrogate the genes with an eye on their roles in fundamental processes at the molecular or channel level, or perhaps at the systems level or at the level of tissues, *etc.*, not as a role of the gene in isolation but rather as its role with respect to interactions with other genes. Regardless of the framework of choice, addressing possible differentiation between psychiatric and neurological disorders through genes’ dynamic interactions and their expressions on brain and bodily tissues critical for the person’s functioning may have new utility to aid in developing targeted treatments. Such treatments may be precisely aimed at mitigating such adverse effects on the brain and on the control of the movements that make up the behaviors examined by these observational diagnoses in the first place.

In this paper we leverage recent advances in the modeling of neurodevelopmental stages involving human embryonic stem cells (hESC), which have made transcriptome data from early development available to the scientific community. Such sharing of data from cultures validated by primary developing tissue offers new opportunities to advance analytical and visualization tools that can potentially facilitate the study of the dynamics of cell differentiation across multiple developmental stages. It can help us shed light on the question of differentiating pools of genes associated with disorders of the nervous systems that may or may not rise to the level of mental illness.

An example of such open access work is by Yao et al. [15], which modeled the early stages of human brain development, including early regional patterning and lineage specification. These authors described cell characterizations amenable to providing benchmark datasets to advance our understanding of the origins of disease of the human brain. Here we use their data to design new visualization tools inclusive of all genes’ fates in the transcriptome and genes’ states across differentiation of self-emerging patterns recorded several times over 54 days. The results available from single-cell RNA sequencing (scRNA-seq) or single cell transcriptomics offer gene-expression data from tens of thousands of genes across hundreds of cells evolving and differentiating towards neuronal stages. These data, combined with identification of disease-associated genes and their expression on human brain and bodily tissues, may help us track the origins of differentiation between psychiatric and neurological disorders.

Analyses of such data often entails dimensionality reduction and visualization of the reduced set (*e.g.,* after Principal Component Analyses, PCA initialization and t-distributed stochastic neighbor embedding, *t-SNE* [16]). Often, during several of the steps leading to the embedding and visualization in much lower dimensional spaces (of two or three dimensions), thousands of genes may be discarded owing to low variability and/or asynchronous expression across various reading days. These data that are discarded may, however, be key to cases where atypical development takes place. Consequently, genes that may be critical to early development toward neuronal stages could be potentially disregarded by current popular methods like t-SNE. This approach would miss an opportunity to examine the transcriptome data from the vantage point of inter-related nodes in a network, using a stochastic approach that goes beyond locally selected neighboring interactions of genes with systematic variability to leverage (and understand) the dynamic, asynchronous nature of many otherwise discarded genes during pluripotent neuronal differentiation.

We propose new methodologies (Figure 1B) that treat a set of genes as a network entity whose parts interact with each other over the course of cell development. To that end, we use a layered approach. First, we identified genes associated with a plurality of psychiatric and neurological disorders, and which also overlapped, thus being associated with phenotypes that are considered comorbid with, for example, autism spectrum disorders (ASD), attention deficit hyperactivity disorder (ADHD), cerebral palsy (CP), *etc*. [9; 10]. Then, we considered the cumulative expression of the genes across four readings through 54 days, as the cells transitioned to neuronal classes. For each gene, we derived marginal distributions of expression across cells and tracked pairwise dependencies to interrogate the full transcriptome dynamically on each reading day. We did so within another layer of inquiry, as the genes formed part of a probabilistic interconnected graph.

This network of interacting interconnected genes associated with a plurality of psychiatric and neurological disorders cannot be *separated* into a disjointed collection of data points, since the topological properties of the graph determine expression and differentiation. We therefore found relationships between local and global properties of this network by defining metrics that quantified the “fate” of each gene during cell differentiation and the degree of interdependency between all cells. Furthermore, our approach was dynamic, *i.e.,* we looked at gene expressions on multiple days for a culture of cells. This approach allowed us to shed some light on how changes in the “state” of the expressions of different genes during the embryonic stages determined the clinical phenotype and characterized different pathologies of the CNS.

This characterization based on the fate and state of the genes’ expressions led to the proposition of a general mechanism and new paradigm that traces the origins of differentiation and commonalities between neurological and psychiatric disorders back to the early stages of embryonic development. This is the point when cells differentiate and become fully matured neurons that will make up the different systems of the nervous system. In this sense, we transformed the current psychiatric *vs.* neurological disorders debate into an opportunity to explore when, during these early embryonic stages, the genes expressed as one disorder or another as a function of their degree of interdependencies. We discuss the implications of our results while considering the notion that gross data such as low-variability and asynchronous genes expression, which are often discarded as superfluous, may in fact hold the key to many unknown aspects of neurological and/or psychiatric disorders. Developing new methods to harvest and utilize their dynamically changing stochastic activities may be critical to understanding the mechanisms guiding us in the design of treatments to cure of diseases.

## 4 Datasets and Codes

Single-cell RNA-seq data during neural differentiation of hESCs were provided by the study of Yao et al [15]. They revealed a multitude of neural cell classes with a range of early brain regional identities. They analyzed 2,684 cells with >20,000 transcripts, as assessed by unique molecular identifiers (UMIs). Cell types were named by point of origin as progenitor (P), transitional (T), neuronal (N) or other tissue (O). We focussed on the evolution of 24,046 genes’ expression across the neuronal type examined at days 12, 19, 40 and 54.

The code that implemented the Kernel Statistical Test of Independence and was used for the present analysis is available in Arthur Gretton et al. [17] as free source material on their website [https://www.gatsby.ucl.ac.uk/~gretton/].

## 5 Methods

*Disorders:* From the genetic database of the Simons Foundation Research Initiative (SFARI) we gathered the set of genes that, according to the literature, are linked to the behavioral diagnosis of Autism. We also compiled several sources found in the DisGeNet portal and identified the lists of genes linked to a variety of neurological (Fragile X-Associated Tremor/Ataxia Syndrome (FXTAS), Dystonia, Cerebral Palsy (CP), Ataxia Syndrome, Tourette’s and Late and Early Onset Parkinson’s disease [18]) and psychiatric disorders [Schizophrenia, Depression, Obsessive Compulsive disorder (OCD), bipolar disorder and Post-Traumatic Stress Disorder (PTSD)]. The psychiatric disorders under consideration are defined in the DSM, whereas the neurological ones are comorbid with autism spectrum disorders (ASD) across the human life span. Some disorders like CP and Tourette’s are also included in the DSM.

*Transcriptome data:* Data matrices expressed as M cells x N genes containing at each entry transcription in log Counts/million. The matrix is transposed to express each of the 24,046 genes as it expresses across cells. This is represented as 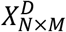, where *N* is the number of genes and *M* the number of cells for day D. Figure 2A shows the counts across neurons at day 40 (top) and at day 54 (bottom) for one gene. Notice that cells may not necessarily be the same on each day, but we work on the genes’ expression space. Next we took the expression of each gene at each cell as peaks in a series (shown in red) and normalized them using equation 1. Notice that the cell order is preserved from the original matrix of cells by genes. It is not a temporal order (it is not a time series), and as such order does not matter in our calculations:

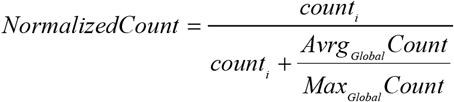

Here *count*_*i*_ is the count value of the *gene*_*i*_ and *Avrg*_*Global*_*Count* is the overall average of the matrix of values taken along the columns and the rows. *Max*_*Global*_*Count* is the maximum count value, also taken globally across the matrix values. We take each such value and apply Eq.1 to scale all expressions of that gene across all cells, each day. The output of the normalization is shown in Figure 2B for cells with expression close to 1. These are cells where the ratio of Avrg_Global_ Count to Max_Global_ Count is very small, so the denominator is only a small margin greater than the numerator. The inset stem plot in Figure 2C (inside the top and bottom histograms) represents all values of the gene across all the cells, scaled as spikes ranging between 0 and 1. These include values for which the ratio of Avrg_Global_ Count to Max_Global_ Count is large and the overall normalized value is small. There is not a temporal order, it is just a series of fluctuations whereby the frequency histogram of all those values (smaller and larger) is of interest.

**Figure 2.**
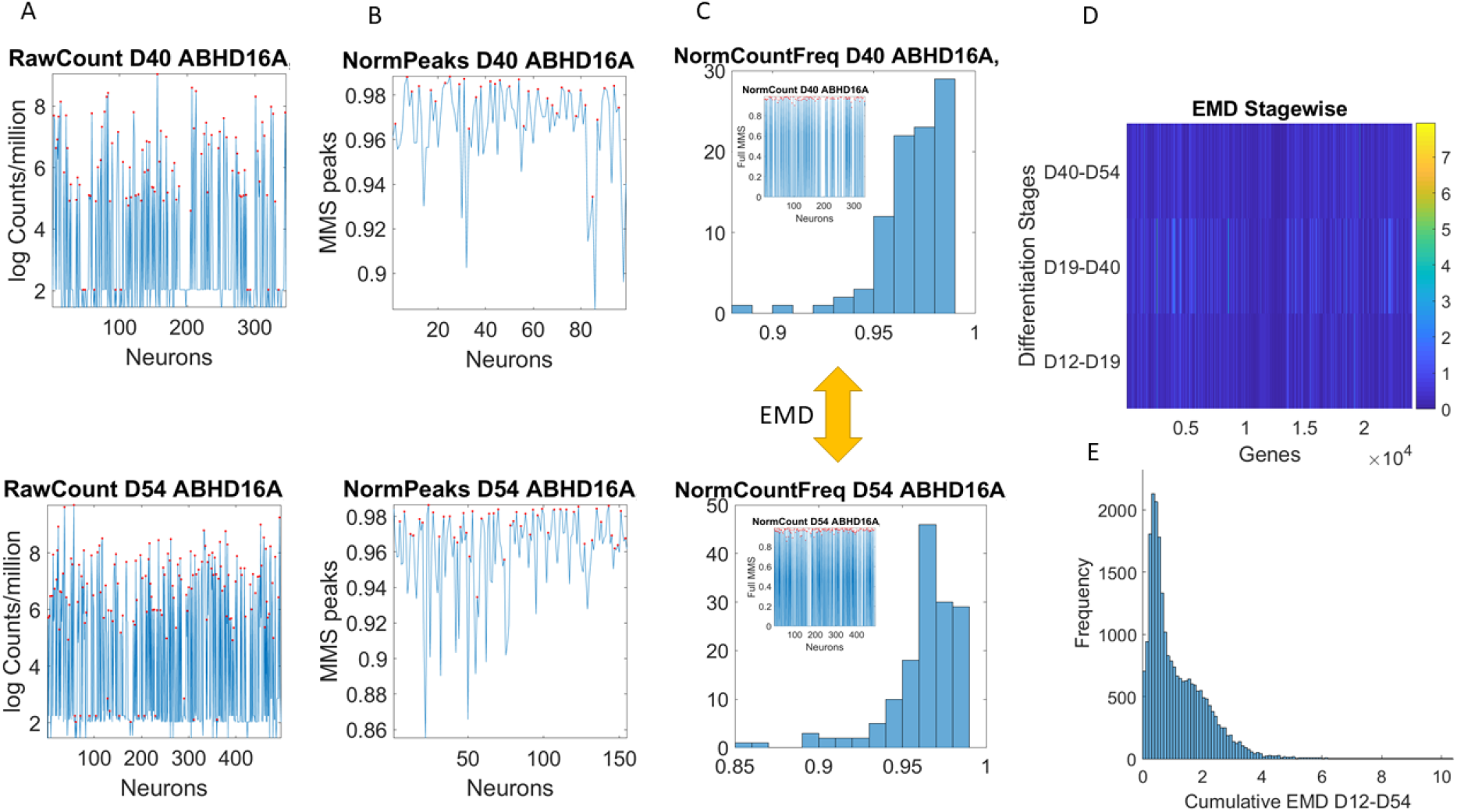
Pipeline to estimate genes’ fates(?) according to expression across cells. (A) Sample gene’s raw count across 496 cells on day 40 and day 54. (B) Normalized gene expression series for range of values obtained using equation 1. (C) Frequency histogram of scaled values across all neurons (represented as spikes between 0-1 in the inset). (D) The earth mover’s distance (EMD is taken between the frequency histogram of consecutive readings (*e.g.,* between the histograms of Day 40 and Day 54) and obtained for each gene (across the columns of the matrix). EMD values range according to the color bar. (E) The cumulative EMD from day 12 to 54 is then obtained for each gene and a frequency histogram of all genes reveals the distribution of all values in fate space. Each bin in this histogram boasts a level of its genes’ expression as they accumulated along the developmental trajectory of the cells. Which genes make up each bin can be easily obtained to further understand genes’ interactions according to fate.

We then obtained the similarity distance between the two frequency histograms corresponding to the gene (across all cells read out each day) using the Earth Mover’s Distance (EMD) [19; 20].

### 5.1 Genes’ Fate: Quantifying it Through the Cumulative EMD

The EMD is a measure of the distance between two probability distributions and is the 1^st^ Wasserstein distance from the family of Wasserstein distance metrics. Essentially, the EMD is the minimal cost of transforming one distribution into the other.

Consider the unknown probability spaces ([0, +∞], 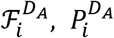) and ([0, +∞], 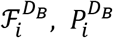) that define the statistical behaviors and corresponding probability distributions for a gene on two different days, *D*_*A*_ and *D*_*B*_, during cell differentiation. The probability distributions can be approximated through histogram fitting from the available normalized cell expression data. Then, *EMD*_*i,A*→*B*_ is the EMD between 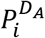 and 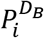 and is indicative of the departure of the statistical behavior of the gene *i* as we move from day *D*_*A*_ to *D*_*B*_. Consider the days 12, 19, 40 and 54 of hESC differentiation. Then we define the quantity:

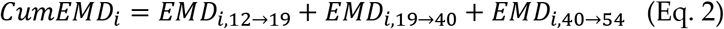

as the cumulative EMD distance of the gene or the “fate” of the role of that gene throughout embryonic stem cell development and differentiation. Then, the average EMD or “fate” of a set of genes *i* = 1, …, *N* associated with a particular pathology is the quantity:

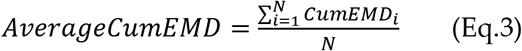

Figure 2D shows a color matrix with all genes across the columns (horizontal axis) and three rows representing, for each gene, the EMD obtained between two consecutive days.

We focused here on graphical modelling techniques and kernel-driven statistical independence hypothesis testing to determine the level at which these genes co-depend [17]. This kernel technique was used to build a parameter space representing 16 disorders sorted out according to their levels of statistical co-dependency at the earliest and latest stages of the cell evolution. Our data of interest were the cumulative EMD traveled representing the gene’s fate.

### 5.2 Kernel Statistical Test to Determine the Level of Interdependence Between Genes Associated with Each Disorder at Different Times During Cell Maturation

On a specific day during cell maturation, for any two genes A and B we had available their expression levels in N cells. We wanted to test whether the expression of gene A was statistically independent from the expression of gene B. Before we present the solution to this problem, let us first introduce the general framework for measuring independence, based on cross-covariance operators in *Reproducing Kernel Hilbert Spaces* (RKHSs).

#### 5.2.1 Cross-covariance operator and Hilbert-Schmidt Independence Criterion

Different methods for measuring statistical independence between random vectors have been proposed over the past decades. Non-parametric approaches to this problem can be traced back to 1948 by Hoeffding [21], when he proposed a test statistic that depended on the rank order of the identically and independently distributed sample data. Techniques that constructed statistics based on empirical characteristic functions were later developed. Modern methods introduced the concept of Kernel Independence Measures, which have found applications in Independent Component Analysis [22; 23], fitting graphical models and feature selection. However, such measures do not necessarily ensure statistical significance. Hence, we decided to use a Kernel Statistical Test of Independence that was developed by Gretton et al. [17], which allowed us to perform hypothesis testing on whether two datasets were independent, and which we applied on the gene expression dependence problem. We briefly present their methodology in the next few paragraphs.

Consider the Hilbert space *F* of functions from a measurable space *X* → *R*. To each point *x* ∈ *X* there corresponds an element *φ*(*x*) ∈ *F* such that < *φ*(*x*), *φ*(*x*′) >_*F*_ = *k*(*x*, *x*′) where *k*: *X* × *X* → *R* is a positive definite kernel. If we assume that *F* is separable, then *F* is a RKHS.

Note, that *F* is the completion of the set of all functions that are linear combinations of these feature functions. To evaluate the value of any function *f* ∈ *F* at some point *x* ∈ *X* one can simply take the inner product between the function *f* and the feature function *φ*(*x*) mapping of the point *x*. This is known as the Reproducing Property, hence the term Reproducing Kernel Hilbert Space. Similarly, we define a RKHS space *G* of functions from a measurable space *Y* → *R* with feature map *ψ*(.) and kernel *l*(.).

Now, assume the probability spaces (*X*, 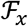, *P*_*x*_) and (*Y*, 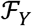, *P*_*x*_) where 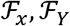 are the Borel σ-fields on *X*, *Y*, respectively, and *P*_*x*_, *P*_*y*_ the corresponding probability measures. Then, for any two functions *f* ∈ *F* and *g* ∈ *G* the cross-covariance operator *C*_*xy*_: *G* → *F* is defined as:

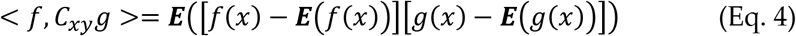

Or with respect to the feature mappings *φ*(*x*), *ψ*(*x*) :

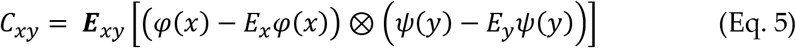

It can be shown that *X* and *Y* are independent if and only if the largest singular value of the operator *C*_*xy*_ is zero. As a measure of independence, the authors consider the Hilbert-Schmidt norm (*i.e.,* the Hilbert-Schmidt Independent Criterion, HSIC) of *C*_*xy*_, which is equal to the sum of squared singular values of *C*_*xy*_ and has a population expression [17]:

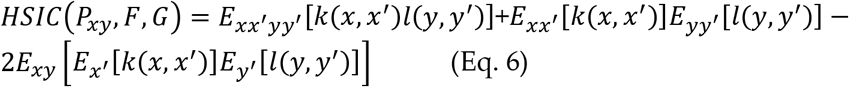

where *x*′ denotes an independent copy of *x* and *k(.)* and *l(.)* are the kernels previously defined. The authors derive an empirical estimate of this independence criterion that follows the V-statistics and has expression:

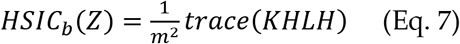

where *Z* is a sample of (*x*, *y*) pairs drawn independently from the distribution of *X* × *Y*, with size *m*, *K* is the *m* × *m* matrix with entries *k*_*ij*_, L is the *m* × *m* matrix with entries *l*_*ij*_ and 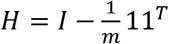, where 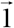 is a row vector of ones.

Then, they proceed to construct a statistical hypothesis testing protocol to test whether *X* is independent of *Y* based on samples (*x*, *y*)^*m*^ drawn from the probability space (*X* × *Y*, 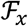 × 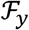, *P*_*xy*_). The null hypothesis is *H*_0_: *P*_*xy*_ = *P*_*x*_ *P*_*y*_ and the alternative hypothesis *H*_1_: *P*_*x*_*y* ≠ *P*_*x*_ *P*_*y*_

Finally, they approximate the independence criterion with a gamma distribution:

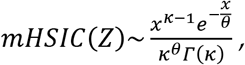

where

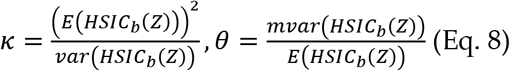

If *mHSIC*(*Z*) is above the threshold determined by the level of significance that we choose for the test, the null hypothesis is rejected. Figure 3 depicts this pipeline.

**Figure 3:**
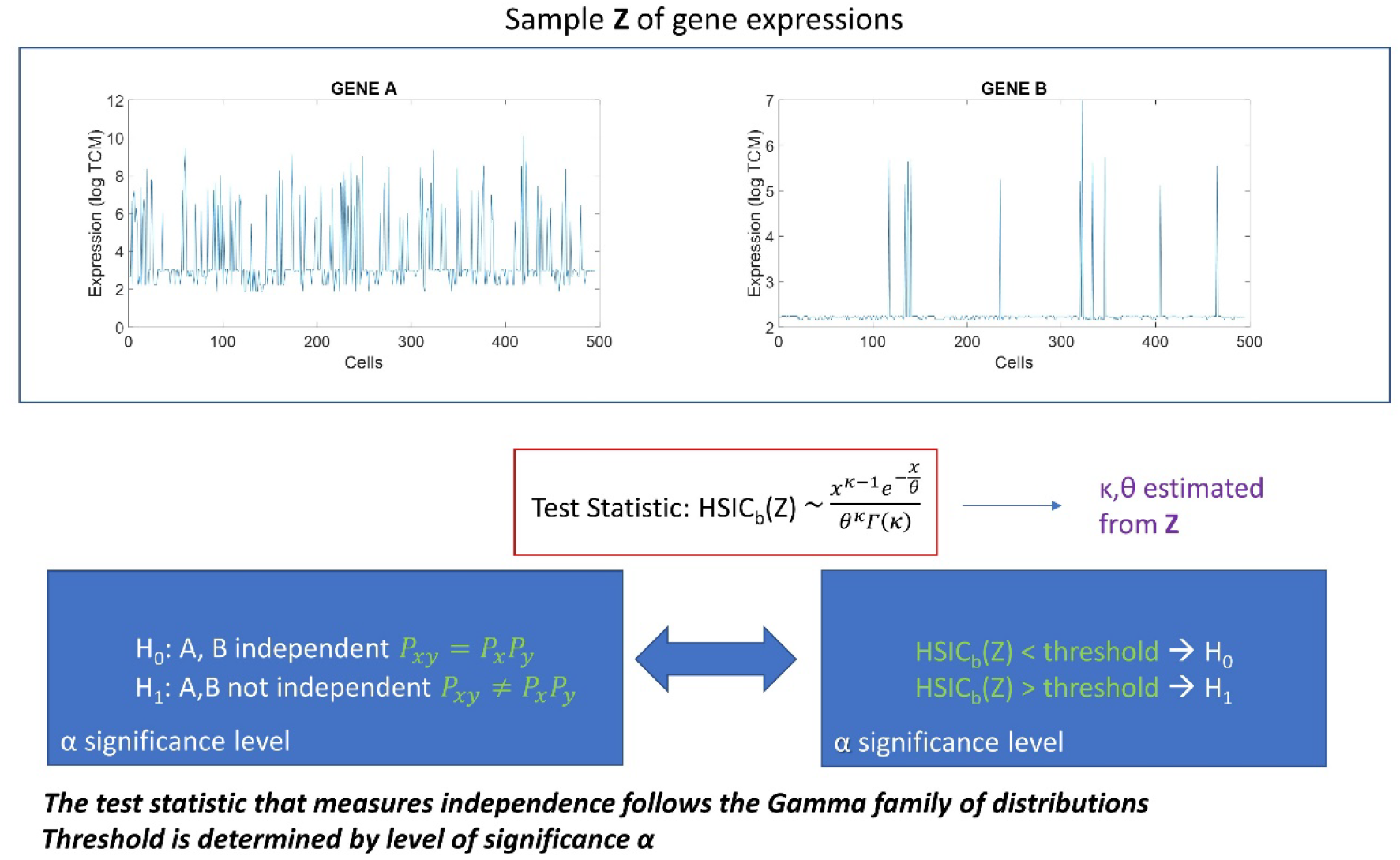
The Kernel Statistical Test of Independence by Gretton et al. [17] applied on the gene expressions of a pair of genes in the hESCs.

#### 5.2.2 The construction of gene expression statistical dependency networks for days 12, 19, 40 and 54 of neural cell maturation

For each disorder, we have a set of genes associated with it (extracted from DisGeNet.) On a particular day D of cell maturation, we have the data 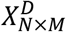, where *N* is the number of genes and *M* is the number of cells. Each row of our data refers to a specific gene and each column to a specific cell.

Define the unknown probability spaces ([0, +∞], 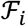, *P*_*i*_) and ([0, +∞], 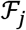, *P*_*j*_) for the expressions of any two genes *i* and *j*, (*i*, *j* = 1, …, *N*) and the probability space ([0, +∞]^2^, 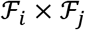, *P*_*ij*_ (for the joint expression of the two genes. Here, [0, +∞] and [0, +∞]^2^ are the measurable spaces for the gene expressions and joint gene expressions, respectively, 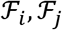 the Borel σ-fields generated by the measurable spaces of gene expressions of *i* and *j*, *P*_*i*_, *P*_*j*_ the corresponding probability measures for the two genes and *P*_*ij*_ the joint *probability* measure. Consider the sample 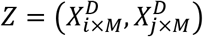, *i.e.,* the pairs of the two gene expressions in the cells. Choosing a level *g* of statistical significance, we can apply the Kernel method on the sample Z and determine whether the genes *i* and *j* are independent of each other on that specific day.

We perform the independency test on all undirected pairs (*i*, *j*), *i* = 1 … . *N* − 1, *j* = *i*, *i* + 1, …, *N* of genes and we construct the graph:

*G* = (*V*, *E*), *V is the set of nodes and E the set of edges*, |*V*| = *N*, *where* |. | *denotes the cardinality of a set. An edge belongs to the graph*, *i.e., e*_*ij*_ ∈ *E if and only if the genes i and j are statistically dependent according to the kernel independency test.*

If the graph were fully connected it would have number of edges 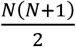, and in this case all genes would be fully dependent upon one another. In search of a metric that shows how interconnected the genes related to the disorder of interest are, we define the *dependency index*:

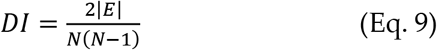

Therefore, for a specific disorder *d*, by constructing the dependency graphs 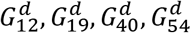 for each day of the disorder we can derive the dependency indexes 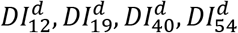. Furthermore, we define:

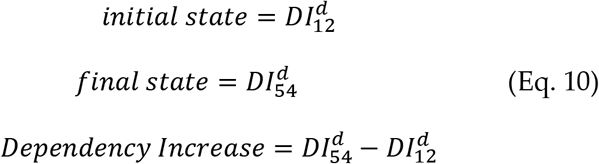

Then, we can map each disorder to a parameter space of dependencies throughout the course of the cell development to track how the interdependence between the genes associated with each disorder evolves through time, from the state of pluripotency to the state of full neural maturation. This allows us to stratify the spectrum of neurological and psychiatric disorders with regards to the complexity of their genotypical expressions.

Using the database DisGeNet, we identified the genes that are associated with each of the following disorders: Fragile X-Associated Tremor/Ataxia Syndrome (FXTAS), Dystonia, Cerebral Palsy, Ataxia Syndrome, Tourette’s, Late Onset Parkinson’s disease (Late PD) and Early Onset Parkinson’s disease (Early PD) and Schizophrenia, Depression, Obsessive Compulsive disorder (OCD), bipolar disorder and Post-Traumatic Stress Disorder (PTSD).

For days 12, 19, 40 and 54 of the hESC differentiation to neural cells, we had the expressions of 24,046 genes from the human genome in log TCM (log transcripts count per million). For days 12, 19, 40 and 54, we had available the expression of all those genes in 263, 168, 346 and 495 different cells, respectively. Therefore, for the sets of genes associated with each of the disorders of interest, we had the datasets 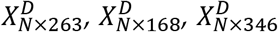, and 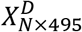 for the four days, where D denotes the disorder of interest and N the number of genes associated with it. From these datasets we calculated the Dependency Indexes (see Methods section) for each day.)

##### Undirected graphical modelling for evaluating the dependency between gene expressions

Graphical models have been extensively researched and used to describe statistical dependencies between random variables. These models can be either undirected graphs or directed graphs; in the latter case, we could derive cause-effect relationships between the variables. In the current project we were interested in undirected graphical models.

Formally, for any set of random variables *X* = (*X*_1_, *X*_2_, …, *X*_*N*_), a graphical model attempts to associate the joint random vector *X* drawn from the probability space (*Ω*_1_ × *Ω*_2_ × … × *Ω*_*N*_, *F*_1_ × *F*_2_ × … × *F*_*N*_, *P*_*x*_) with a graph *G* = (*V*, *E*), where *V* stands for vertices and *E* for edges. Here, *Ω*_1_, *Ω*_2_, …, *Ω*_*N*_ are the sample spaces for each random variable and *Ω*_1_ × *Ω*_2_ × … *Ω*_*N*_ the joint space of the random vector *X*. *F*_1_, *F*_2_, …, *F*_*N*_ are the corresponding generated Borel σ-fields to denote the sets of all possible random events for each random variable and *P*_*X*_ is the probability measure on the random vector *X*. The set of nodes *V* represents the random variables *X*_1_, *X*_2_, …, *X*_*N*_ drawn from the probability spaces (*Ω*_1_, *F*_1_, 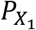), (*Ω*_2_, *F*_2_, 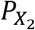), … (*Ω*_*N*_, *F*_*N*_, 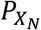), with their respective Borel σ-fields and probability measures. An edge *e*_*ij*_ ∈ *E* if and only if the random variables *X*_*i*_ and *X*_*j*_ depend on each other.

If two nodes *u*_*i*_ and *m*_*j*_ are not connected with an edge it implies that the two variables *X*_*i*_ and *X*_*j*_ are conditionally independent, *i.e*., statistically independent *given* all other nodes:

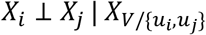

This property of the graphical model is known as the *global Markov property*.

#### 5.3.1 General estimation of a graphical model using Chow-Liu trees

If we want to factorize the joint probability distribution in a dependency graph that has a tree structure, *i.e.*, every two nodes are connected by no more than one path (in other words there are no loops in the graph), then the joint density of the random vector X factorizes with respect to the pair-wise joint and marginal densities as:

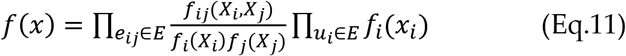

It turns out that, in the case in which all variables are categorical and take values from a finite set, it is very easy to find the optimal tree that factorizes the joint distribution. Let *N*_*x*_ be the number of times a realization *x* of the random vector *X* appears in a collection of independent and identically distributed (*i.i.d.*) samples. The tree *G* that optimally factorizes *X*, given the sample data *Z* of size *n*, will be the one with the maximum log-likelihood (MLE):

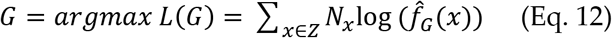

which turns out to be:

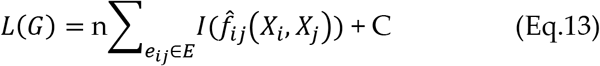

where C is a constant and 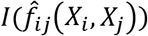 the empirical mutual information between *X*_*i*_ and *X*_*j*_. Therefore, by choosing the appropriate subgraph G that maximizes the sum of the empirical mutual information estimates, we obtained the optimal tree structured graphical model. Since the model is a tree, we simply needed to find the Maximum Spanning Tree from the mutual information network, for example, by using Kruskal’s algorithm. The process we just described is known as the Chow-Liu algorithm and the extracted conditional dependency tree that factorizes *X* is the Chow-Liu tree.

#### 5.3.2 Gene’s co-dependencies graphical models spanning (Chow-Liu) trees

For a specific set of diseases, in this case neurological and psychiatric, we had a set of genes associated with them. On a particular day D of cell maturation, we had the data 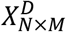, where *N* is the number of genes and *M* the number of cells. Each row of our data referred to a specific gene and each column to a specific cell. We treated each column (corresponding to a cell) as an *i.i.d.* sample drawn from the joint probability space of the expressions of that set of genes, and we wanted to generate the undirected graphical networks *G*_12_, *G*_19_, *G*_40_, *G*_54_ for days 12, 19, 40 and 54.

In the case of continuous variables, the methods used to estimate the Chow-Liu tree usually involves Kernel Density Estimation (KDE) of the joint and marginal probability densities [24]. However, in our case we wanted to factorize the joint probability density of a gene expression network with number of cells on the order of magnitude ~10^2^. The application of KDE, given the dimensionality of the data (number of genes), would require in this case a number of samples far exceeding the available number of ESCs. Therefore, we resorted to extracting the Chow-Liu Trees by estimating the mutual information through binning and histograms on the available data [25]. Once the Chow-Liu Tree corresponding to the factorization of the gene set of interest was obtained, we ordered the nodes in ascending degree. The proposed methodology can be appreciated in Figure 5 of the methods section.

**Figure 4.**
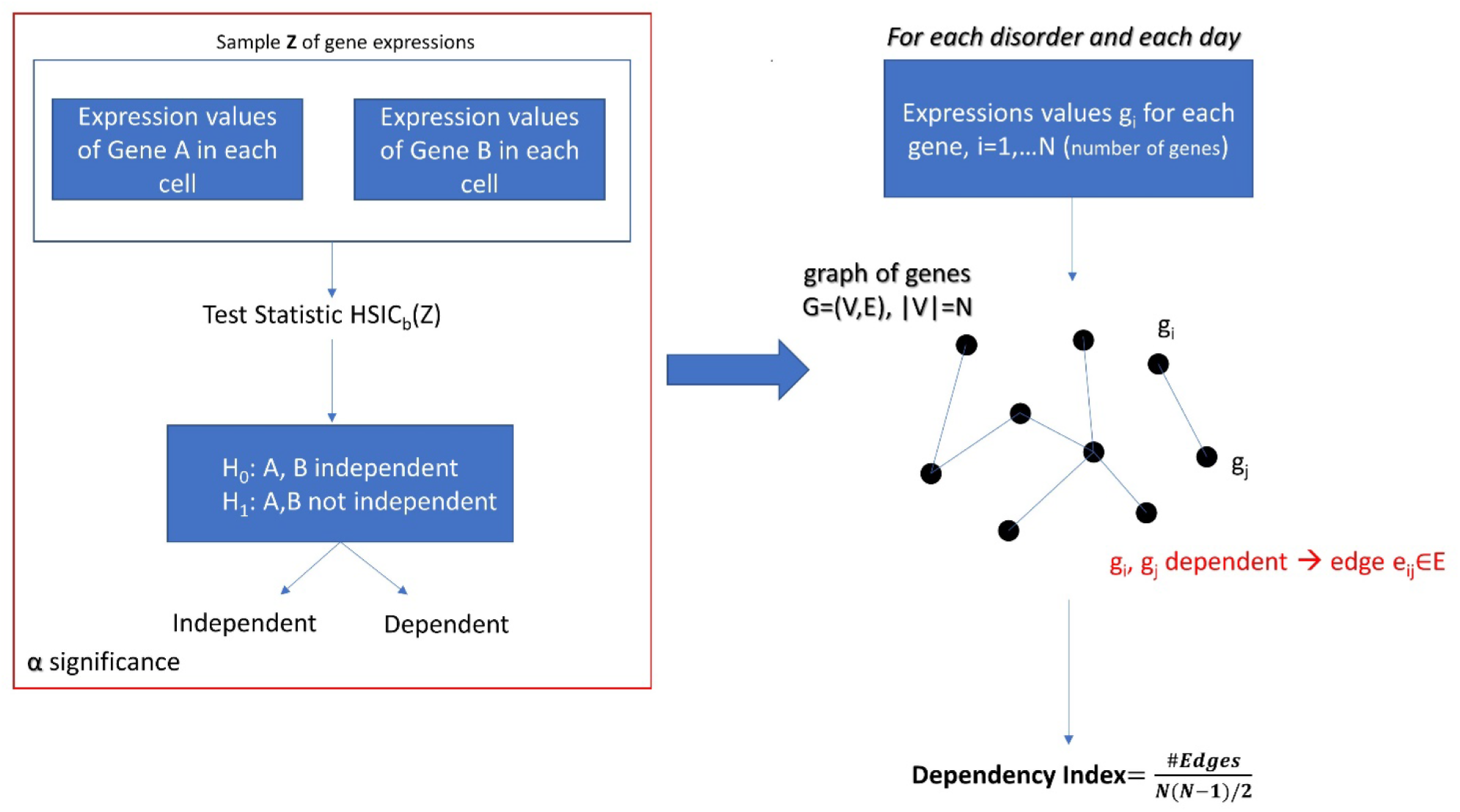
The construction of a graphical model after performing the Kernel Statistical Independency Test on every pair of gene expressions associated with a particular disorder on a specific day of cell differentiation. The Dependency Index characterizes the degree of interdependence of genes that define the nodes of the graph.

**Figure 5:**
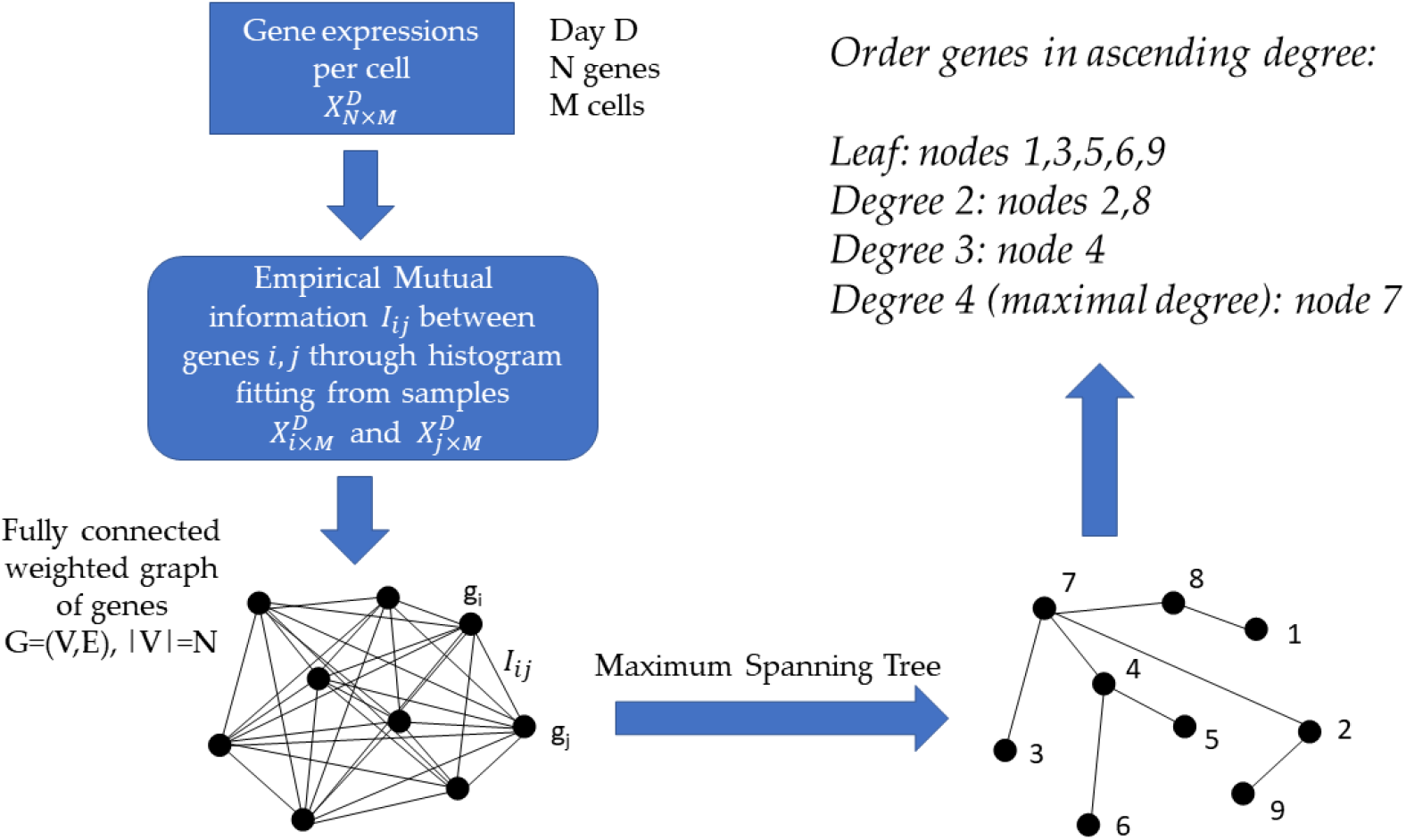
Proposed pipeline for the factorization of the joint probabilistic behavior of a network of genes and for determining the significance of each gene in the network

What do we achieve with this ordering? It is obvious that the higher the degree of a gene is, the more co-dependent its expression is with the expression of other genes in the network. This implies, in a statistical sense, that mutation or deletion of this gene is bound to immediately affect the expressions of many other genes. Note that this rationale simply states the co-dependency between the gene expressions, but the actual (causal) mechanisms through which this statistical relationship takes place are open to investigation.

#### 5.3.3 The gene’s state through a binary code

We obtained the average degree across the network’s nodes each day. We then set it as the threshold value to determine ON or OFF state for the gene each day. Across 4 days, we had 16 possible binary states (2^4^) that provided the state trajectory of the gene (in addition to its fate.) This information served to classify cells according to the genes type. To that end, for each cell, we obtained the frequency histogram of counts corresponding to each class of genes, [0000],[0001],…,[1111]. Then, we used MLE to obtain the probability distribution best characterizing the histogram and obtained a parameter space to represent the shape and scale parameters thus determined as points along a scatter. Since each gene has a binary configuration, for a given day, we could then retake the cells x genes matrix and ask which cells cluster in this space according to the 16 possible states.

## 6 Results

### 6.1 Cumulative EMD Captures Dynamic Evolution of the Genes’ Expression in Fate Space

The contributions of each gene to the overall evolution of the hESCs as they reached neuronal stages were well captured by the stochastic characterization of their normalized count taken for each gene across cells each day. These marginal distributions can be seen evolving across all genes in Figure 6A, which shows the frequency histogram of all EMD values obtained for each comparison. Figure 6B shows the values sorted out across the genes, providing a sense of the overall trajectory of changes in gene expression across all cells. Also notice that, since each day the number of cells changes, we normalized the EMD quantity to range between 0 and 1 when superimposing the data across days (Figure 6B.)

**Figure 6.**
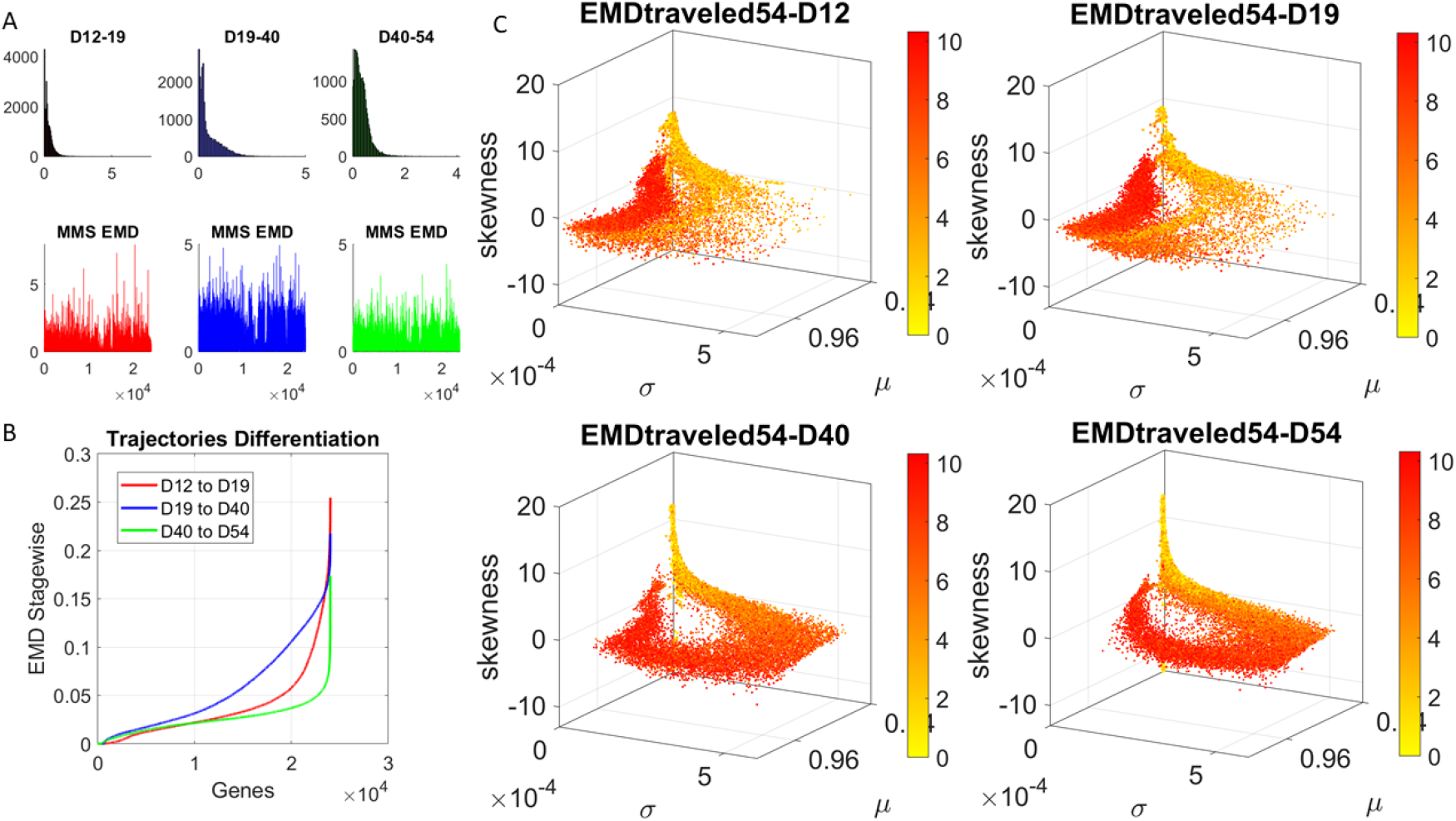
Dynamic gene’s motion trajectory in fate space. (A) The distributions of EMD obtained for each measurement between days (top frequency histograms) for all 24,046 genes and bottom values across genes each day. EMD based on normalized counts using Eq.1. (B) Sorted values of change in EMD across genes showing the trajectory from measurement to measurement, for each gene (horizontal axis). Order is based on the cumulative EMD at day 54; then we plotted the values of EMD at each stage (D12-D19, D19-D40, D40-D54). (C) Full transcriptome representation on fate space is obtained by representing each point as the empirically estimated Gamma PDF moments for the best fitting shape and scale parameters (in an MLE sense). The mean μ is represented along the *x-axis*, the variance σ along the y-axis and the skewness along the *z-axis*. The kurtosis is proportional to the marker size (larger represents more kurtotic distribution). Color bar reflects the value of the cumulative EMD at day 54 when the cells are neurons. Then, we trace back where the gene was in fate space at the initial stages, D12, D19 and D40.

We then accumulated the EMD value by summing up to day 54 and building a colormap to visualize the change across all genes in the transcriptome. For each gene, we took the frequency histogram of the normalized gene expression across the cells and, using maximum likelihood estimation (MLE), we obtained the continuous family of probability distributions that best fit the histogram in an MLE sense. This was the Gamma family, which spans a broad range of shape (*a*) and scale (*b*) values (ranging from the memoryless exponential *a* = 1, to distributions with heavy tails, to symmetric, Gaussian-like distributions.) We estimated the Gamma moments of the empirical distribution corresponding to each gene. Each day we plotted them on a parameter space, whereby the mean is represented along the *x-axis*, the variance is represented along the *y-axis*, the skewness is represented along the *z-axis* and the kurtosis is proportional to the size of the marker (higher kurtosis represented by larger marker size.) We then colored each gene with the cumulative EMD at day 54, when the cells had reached neuronal state. This enabled us to visualize the ‘motion’ dynamics of the 24,046 genes in the transcriptome and, retrospectively, to see where the most active and least active genes were located. This information is shown in Figure 3C, as the cells evolved to neuronal stages.

### 6.2 Different Evolution of the Dependency Index Values Throughout hESC Maturation for Psychiatric *vs.* Neurological Disorders

We obtained the quantity Dependency Increase, as explained in the methods, by taking the shift in the Dependency Index at Day 12 versus the increase in the Dependency Index until full neuronal maturation at day 54. We then plotted that value for each disorder along the *y-axis* of a parameter space as a function of the Dependency Index at the initial stage and at the final stage. This is represented as a vector field (rate of change) in Figure 7A. A pattern emerged across disorders whereby all disorders that were classified as neurological, except for Late Onset Parkinson’s Disease, tended to be characterized by a lower dependency during the first stage of cell differentiation and a high increase in dependency at the neuronal stage. The genes associated with these neurological disorders tended to increase their co-dependencies as the cells matured into neurons, but early in the process they were less co-dependent (lower values of Dependency Index on the *x-axis*). These results can be appreciated in Figure 7B.

**Figure 7.**
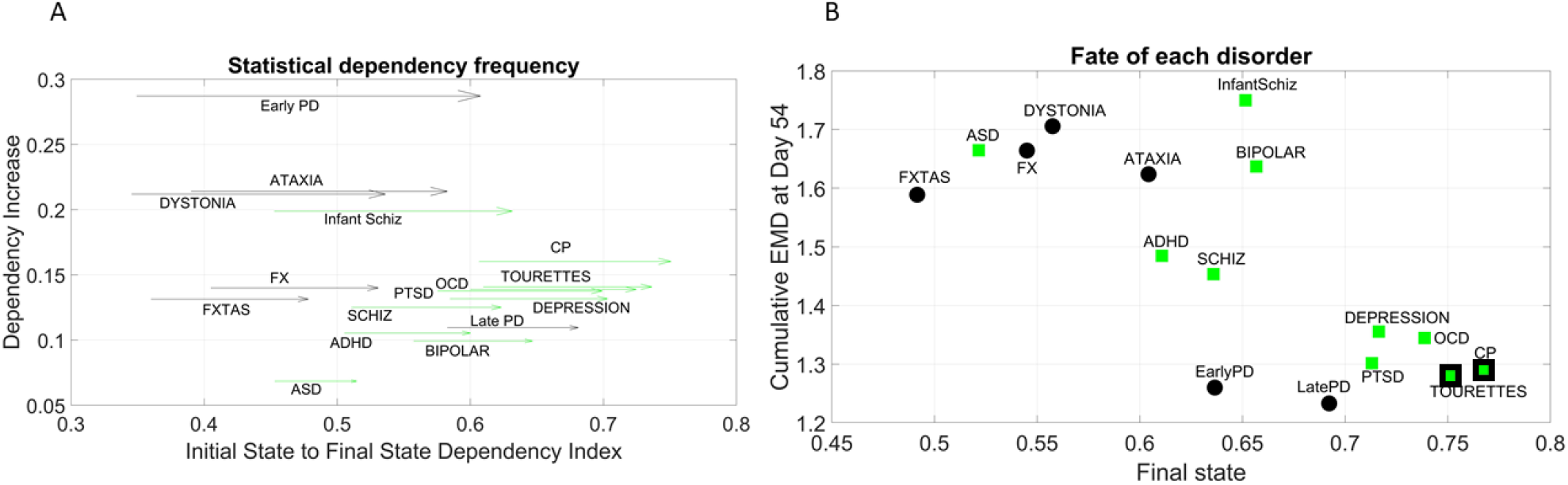
Dynamic evolution of genes’ co-dependencies in neurological *vs.* psychiatric disorders. (A) A negative trend is observed between Dependency Increase and Dependency Index value at the initial state. Psychiatric (green squares) disorders tend to cluster away from neurological (black circles) disorders and exhibit a high degree of dependency in the initial state. ASD lies on the border between most neurological and psychiatric disorders with regards to the dependency metric. Late PD lies among the psychiatric (DSM) disorders. Vector represents the change from Day 12 to Day 54. (B) The final fate of the genes associated with various disorders is plotted as a function of the interdependency index at the final state. As days progress, genes with a high variability in their expression profile (active genes) tend to have a lower interdependency whereas those with a lower variability (lazy genes) tend to have a higher interdependency. Disorders diagnosed by the DSM (green squares) as psychiatric tend to cluster towards high interdependency and lower variability and level of expression. These include depression, OCD, CP, Tourette’s, and PTSD, which group with late and early PD (neurological disorders.) CP and Tourette’s are also considered neurological by some physicians, so the edge of the marker is colored black to represent the mix. At the other end, neurological disorders (FX, FXTAS, Dystonia and Ataxia) cluster with ASD (a DSM disorder with neurological underpinnings.) ADHD is separable from ASD in this parameter space, despite their allowed comorbidity by the DSM-5. ADHD is closer to schizophrenia, whereas bipolar and infantile schizophrenia stand on their own in the middle of the graph and away from the general trend. These disorders have associated genes with mid-level of interdependencies and high EMD at fate, signifying higher variability in expression profile.

In contrast, disorders classified as psychiatric in the DSM-5, as well as Late Onset Parkinson’s Disease, tended to be characterized by a higher dependency during the first stage (higher values along the *x-axis*) but a lower increase in dependency as the cells matured into neurons. Tourette’s, a psychiatric disorder according to the DSM-5, was also traditionally classified as neurological and often labeled as ASD. We highlighted this duality with the black edge of the marker in CP and Tourette’s. In this parameter space, ASD seemed to lie on the border between neurological and neuropsychiatric disorders, whereas infantile schizophrenia (which used to be defined as autism at some point) lined up with ASD along the early value of the Dependency Index, but with a much higher dependency increase as cells matured into neurons.

#### Negative Correlation Trend Characterizes Genes Associated with Neurological and Psychiatric Disorders, with Complementary Features in ASD and PD Associated Genes

For each disorder, we calculated the average cumulative EMD at Day 54 for the genes associated with the disorder (*y-axis*) and plotted it as a function of the Dependency Index at Day 54 (*x-axis*) in Figure 7B. We noticed a negative trend whereby the variability of expression, as quantified by the cumulative EMD from measurement to measurement, tended to decrease for Depression, PTSD, OCD, CP, Tourette’s (all DSM disorders.) In this cluster, early and late PD, which are neurological disorders, appeared amid psychiatric DSM disorders.

Neurological disorders such as FXTAS, FX, Dystonia and Ataxia were high in cumulative EMD but tended to have lower dependency indexes than DSM disorders. ASD, a DSM psychiatric disorder, appeared among the neurological cluster with a high cumulative EMD at day 54 but a lower dependency index during this final state.

Infantile schizophrenia and bipolar disorder, both DSM disorders, were the exception to this negative trend, as they were both high in EMD expression and dependency index.

#### Visualization in Fate Space of Genes Associated with Psychiatric and Neurological Disorders

Using the visualization in Figure 6C, we tracked in fate space the evolution of the genes associated with the neurological and psychiatric disorders. We found that they moved from a spread-out configuration in earlier days toward more localized regions along a path of high variability in EMD and an opposite region of low variability in EMD. As explained previously, the EMD measured the change from one frequency histogram (marginal distribution) on gene’s expressions across the cells on one day to the frequency histogram on the next day. As such, cumulatively they reflect the overall variability in gene expression towards the cells’ fate. We call those with the higher cumulative EMD *active genes* and those with the lower cumulative EMD *lazy genes*.

By day 54, two distinct lobes are obvious in Figure 8, which we projected in Figure 9 along the plane spanned by the first two empirically estimated Gamma moments, mean *vs.* variance. There we saw the high mixture between the neurological (green) and psychiatric (black) disorders. We then asked if there were distinctions across the genes that we could visualize using the Chow Liu maximal spanning trees, treating their network interactions according to probabilistic graphs.

**Figure 8.**
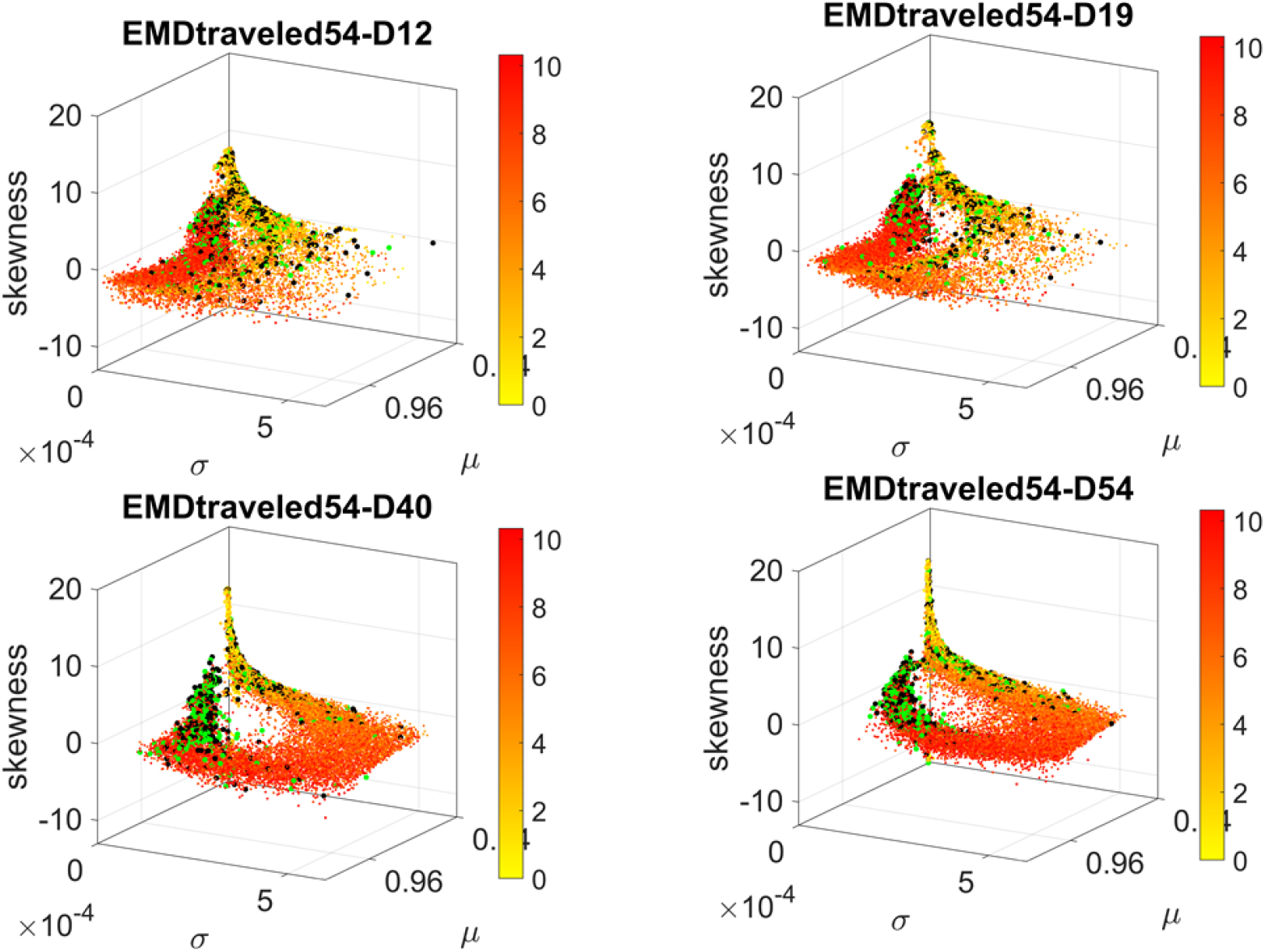
Visualizing the trajectories of genes associated with psychiatric (black) and neurological (green) disorders as they evolve in fate space. Color gradient represents the cumulative EMD at day 54, with yellow representing low values owing to low variability and lower expression whereas red represents high variability and higher expression across cells. At day 54, two lobes emerge along the high values space (higher μ) and central tendency (toward 0 skewness) with higher concentration in lower variance regions (along the σ dimension.) Notice the high change from Day 19 to Day 40, with clearly two lobes with intermixed genes from both classes of disorders in day 54.

**Figure 9.**
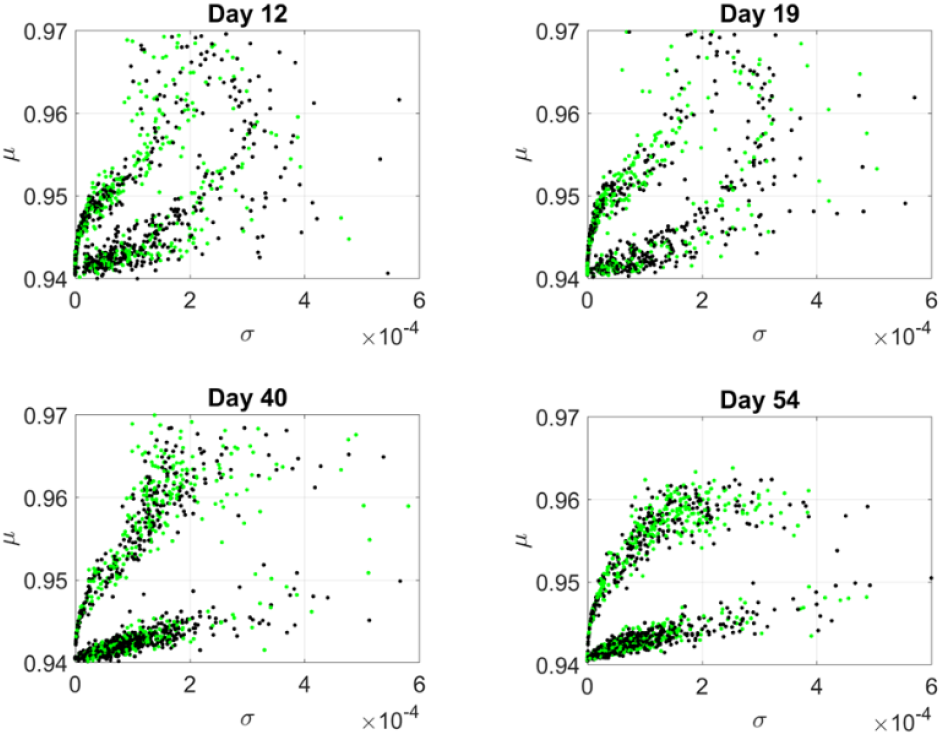
Projection of transcriptome genes associated with neurological (green) and psychiatric (black) disorders on the Gamma moments plane spanned by the empirically estimated Gamma μ and Gamma σ reveals two classes of genes. At each measurement day the genes move to eventually form two distinct lobes corresponding to the active and lazy genes on the curved and line-like scatters, respectively.

The results from the visualization prompted us to further examine these intermixed genes and their evolution along the Gamma mean (μ) and Gamma variance (σ) dimensions of fate space. Projecting these values on a mean, variance parameter plane, clearly showed their evolution and convergence to two lobes in Figure 9 (Day 54), whereby the lobe of active genes with higher variability in EMD quantifying probability transitions separated from the lobe of lazy genes with lower variability. The composition of these lobes was highly intermixed, with 2,613 genes total, 946 in the lower lobe (line like) and 927 in the curved lobe. We then observed 512 unique genes in the curved lobe of active genes and 927 unique genes in the line lobe of lazy genes. These genes might be associated with more than one psychiatric and/or neurological disorder, or just with one or the other (See Appendix Table 1 and expanded Supplementary Material Table). Appendix Figure 1 shows the dependency indexes and fate across disorders split according to genes in these different lobes. We further analyzed the composition of the two lobes, but first, we had a look at the network analyses.

### 6.3 Differentiation of Network Evolution in Active and Lazy Gene Lobes in Psychiatric (DSM) *vs.* Neurological Disorders

The Chow Liu maximal spanning trees for each of the lobes identified in Figures 8 and 9 were obtained and the degree associated with each gene was quantified to represent in Figures 10 and 11 the network evolution of the active and lazy genes, respectively. These graphs show the genes blindly, without the disorders’ labels, to give a sense of the differentiation between the two lobes of intermixed genes in both neurological and psychiatric disorders.

**Figure 10.**
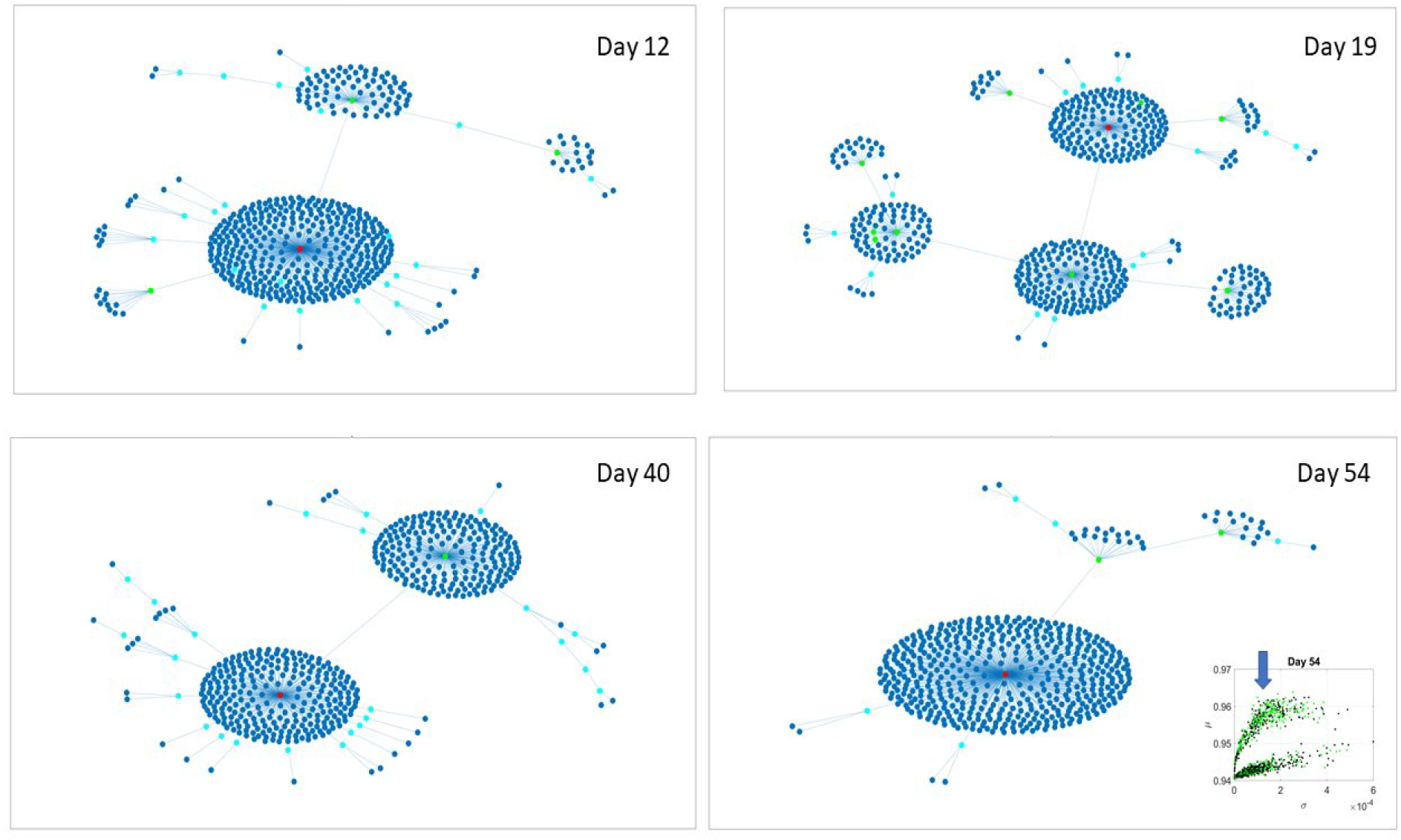
Evolution of statistical dependencies of the active genes. Hubs with two or more nodes but fewer than 10 are cyan; those with more than 10 degrees but not the maximum are colored green. The maximum degree is colored red. In the inset in the lower right panel, the arrow points at the lobe of active genes comprising the dynamically evolving network.

**Figure 11.**
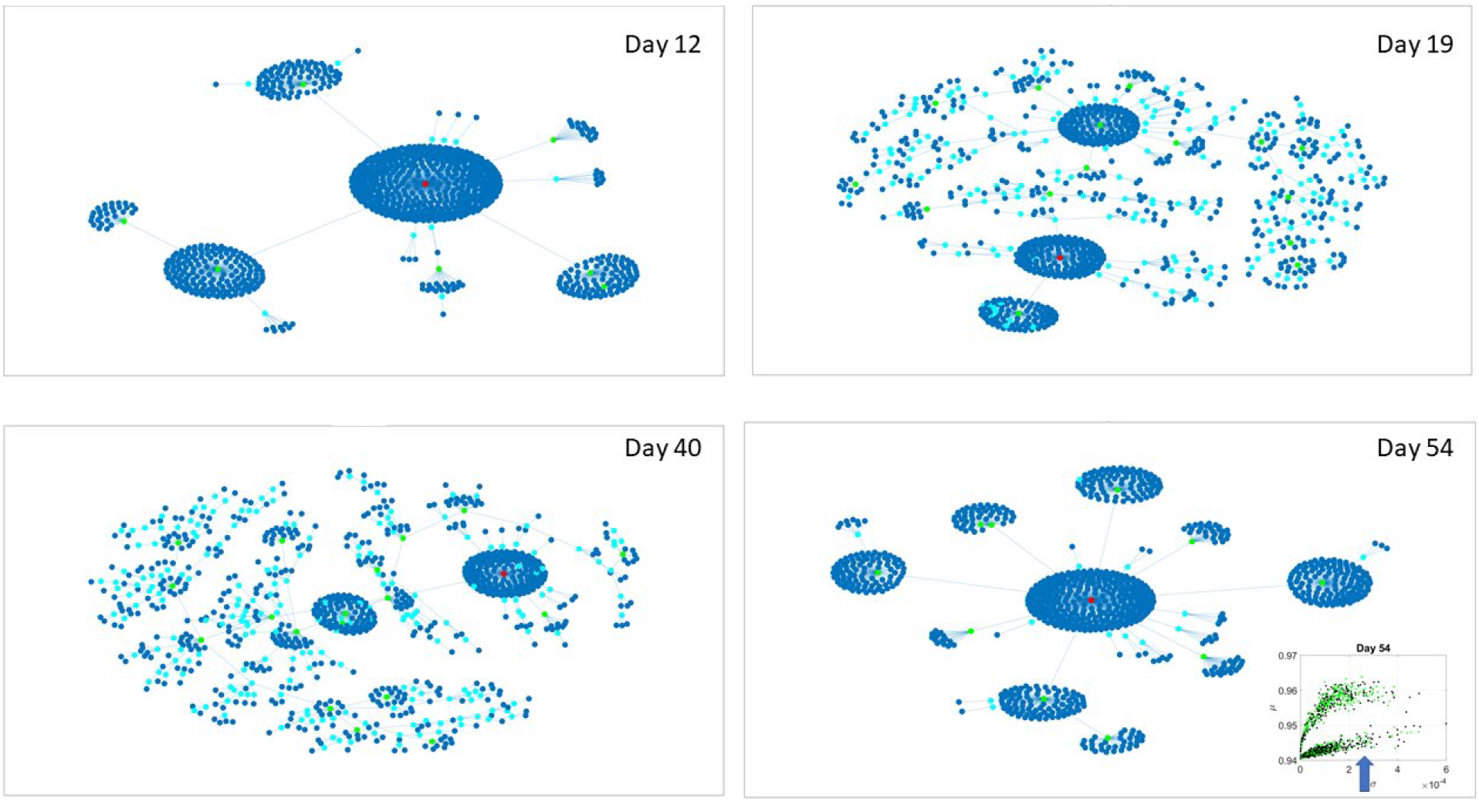
The Chow Liu Maximal Spanning Tree represented as an interconnected network evolving from Day 12 to Day 54, for the lazy genes (located on the line like lobe as marked by the lower right hand inset.) The inset in the lower right panel indicates the lobe of lazy genes comprising these dynamically evolving networks.

Several important conclusions emerge from these representations of the genes’ fate evolution. One is that these two lobes have fundamentally different evolution in genes’ interactions. Another one is that they have a handful of hubs with high co-dependencies among many genes, such that if a link is disrupted between two of these hubs, the network is disconnected with potential cascade effects. Third, days 12 and 54 have fewer hubs than intermediate days. The latter is true in both lobes but more so in the lobe of active genes. The lobe of lazy genes has a far more distributed network in intermediate days 19 and 40, clearly showing the importance of such genes in the overall evolution of the transcriptome toward neuronal states.

These patterns prompted investigation of the hub’s evolution across neurological and psychiatric disorders. We built matrices of hub genes along the rows, sorted by graph degree, and disorders across the columns, sorted by neurological (early PD, late PD, Dystonia, Ataxia, FXTAS, and FX) and psychiatric (DSM diagnosed including infantile Schizophrenia, Schizophrenia, ADHD, ASD, Bipolar, PTSD, CP, Depression, OCD and Tourette’s.) Each entry was color coded with the degree (in log scale) to help visualize the colors better, since they ranged non-linearly from 1 to 470. In the case of degree 1, two genes co-depended on each other’s expression patterns. In the case of degree 2 or more, the node (gene) had an edge with co-dependency with two or more genes. In Figure 11 we focused on the active genes from the curved lobe on the projection of the genes in fate space onto the (μ, σ)-plane (with genes colored in green representing those associated with psychiatric disorders and those colored black associated with neurological disorders.)

We saw that schizophrenia (being the disorder associated with the highest number of genes across these disorders) was the one with the highest number of hubs. We also observed the involvement of a hub across multiple disorders of the neurological class and of the psychiatric class. Furthermore, we noted that psychiatric disorders in general spanned more hubs, and these hubs had a higher degree than neurological disorders. Hubs with degree of 3 and above that were common to both classes of disorders and / or in only one class abounded and are listed in Table 1 of the Appendix and in the expanded Table 1 in Supplementary Material listing also function and other information about the genes.

**Table 1.** Cataloguing active (red) and lazy (yellow) genes by node degree and hub categories by day. See main text Figures 10-11 of evolving networks.

| Neurological and Psychiatric | Genes | Degree | Day |
| --- | --- | --- | --- |
| Ataxia, Schiz, ADHD | **YWHAG** | 7 | 12 |
| FXTAS, SCHIZ, Depression | **YWHAZ** | 78 |  |
| Ataxia, Depression | **ABCC8** | 19 |  |
| Ataxia, FX, Schiz, ADHD, Depression | **APP** | 20 | 19 |
| Ataxia, Schiz | **PGK1** | 45 |  |
| Late PD, Schiz | **ELAVL4** | 165 |  |
| Late PD, Ataxia, Depression, Tourette's | **NLRP3** | 7 |  |
| Dystonia, ADHD, ASD | **CDKL5** | 20 |  |
| Early PD, Late PD, Dystonia, Schiz, ADHD, PTSD, Depression | **SLC18A2** | 20 |  |
| Dystonia, ADHD, ASD | **CDKL5** | 3 | 40 |
| Late PD, Schiz, Depression | **IL2RA** | 3 |  |
| Late PD, FXTAS, Depression | **SGCA** | 3 |  |
| Late PD, Schiz, ADHD, Depression, Tourette's | **SLC6A2** | 3 |  |
| Late PD, Schiz, ADHD, PTSD, OCD | **TAL1** | 3 |  |
| Late PD, Ataxia, Schiz, Depression | **TGFB1** | 3 |  |
| Dystonia, Schiz | **GCHFR** | 4 |  |
| Ataxia, Depression | **NR5A1** | 4 |  |
| Ataxia, Schiz | **PCYT1A** | 4 |  |
| Late PD, ASD | **BRAF** | 7 |  |
| Ataxia, Schiz | **RFT1** | 13 |  |
| Late PD, Schiz | **STH Chr17ctg5 hap** | 26 |  |
| Late PD, Schiz | **WNT2** | 16 | 54 |
| Late PD, Schiz, ADHD, PTSD, Depression, OCD, Tourette's | **IL1B** | 6 |  |
| Ataxia, Schiz | **IL12A** | 7 |  |
| Early PD, Schiz | **CDNF** | 33 |  |
| Late PD, Schiz, ADHD, Depression | **MSMB** | 342 |  |
| Neurological Only |  |  |  |
| None |  |  | 12 |
| None |  |  | 19 |
| Dystonia, Ataxia | **KCNA1** | 6 | 40 |
| Late PD, FXTAS | **SGCA** | 3 | 54 |
| Psychiatric Only |  |  |  |
| Schiz, ADHD, Depression | **SCD** | 11 | 12 |
| Infantile Schiz, ASD | **PXDN** | 5 | 19 |
| Schiz, PTSD, Depression | **STMN1** | 7 |  |
| Schiz, Depression | **DENND5B** | 13 |  |
| Schiz, Depression | **CDKN1C** | 78 |  |
| Schiz, ASD, PTSD | **SET** | 156 |  |
| Schiz, ADHD, Depression | **ADRA2C** | 4 |  |
| ADHD, Depression | **MIRLET7D** | 6 |  |
| Schiz, ASD | CNTNAP3 | 6 |  |
| OCD, Tourette's | POU1F1 | 8 |  |
| Schiz, Depression | CRHBP | 11 |  |
| PTSD, Depression | PKD2L | 12 |  |
| Schiz, Bipolar, Tourette's | ILRL1 | 20 |  |
| Schiz, Depression | MALAT1 | 153 |  |
| Schiz, ADHD | PPIA | 5 | 40 |
| Schiz, Depression | CRHBP | 2 |  |
| Schiz, ADHD, Depression | ADAMTS2 | 3 |  |
| Schiz, PTSD | DUSP2 | 3 |  |
| Schiz, ASD | HS3ST5 | 3 |  |
| Schiz, Bipolar, Tourette's | IL1R1 | 3 |  |
| Schiz, ADHD, PTSD, Depression, OCD | NPSR1 | 3 |  |
| Schiz, ADHD | PRKG1 | 3 |  |
| ADHD, Depression | GALR1 | 4 |  |
| Schiz, Depression | LIF | 4 |  |
| ADHD, Depression | SIM1 | 4 |  |
| ADHD, Depression | CHRNA6 | 5 |  |
| Schiz, ADHD | DGKH | 5 |  |
| Schiz, Bipolar, Depression | NPAS4 | 5 |  |
| Schiz, Depression | TREM1 | 5 |  |
| Schiz, ASD, Depression | RAPGEF4 | 6 |  |
| Schiz, Depression | KLK8 | 7 |  |
| Schiz, OCD, Tourette's | LCT | 7 |  |
| ASD, OCD | CDH9 | 9 |  |
| ADHD, ASD | IQSEC2 | 9 |  |
| Schiz, Depression, OCD | PGC | 9 |  |
| Schiz, ADHD | PIWIL4 | 10 |  |
| Schiz, Depression, OCD, Tourette's | HTR3B | 11 |  |
| Schiz, ADHD | CNTNAP3 | 20 |  |
| Schiz, Depression | MALAT1 | 179 |  |
| Schiz, ADHD, ASD, Depression | TCF4 | 19 | 54 |
| Schiz, ADHD | CD40 | 15 |  |
| ADHD, OCD | GJB2 | 28 |  |

To further investigate the balance between active and lazy genes in neurological *vs*. psychiatric disorders, we quantified the disease ratio,

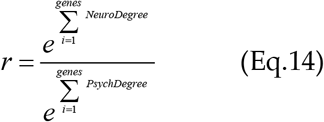

Here the numerator sums over the number of genes with degree above 3 in each of the neurological disorders under consideration and the denominator sums over the number of genes with degree above 3 in each of the psychiatric disorders under consideration. If in a psychiatric disorder (in the denominator) 0 genes participate, the contribution of the denominator is e^0^ = 1. Otherwise, the ratio will reflect the balance of genes in one *vs.* the other, obtained for active and lazy genes separately. Figures 12 (active genes) and 13-14 (lazy genes) depict the matrices whose entry is the scalar value of the ratio (plotted as a color map in logarithmic scale.) Appendix Figure 2 depicts the active genes values and Appendix Figures 3–4 do so for the lazy genes. Figure 12B shows the active genes lobe as depicted on Figures 8-9 for day 54, whereas Figure 12C provides the same plot (rotated) for genes in DisGeNet found on the transcriptome at day 54. In this figure, the size of the marker is proportional to the scalar ratio, reflecting the balance between neurological and psychiatric genes for each class of genes. Figure 12D provides the probability graphs depicting the higher probability for active genes in neurological conditions. This higher proportion comes despite the far larger number of DisGeNet genes associated with psychiatric disorders like Schizophrenia (as captured by the colormap matrices in 12A and 13-14.)

**Figure 12.**
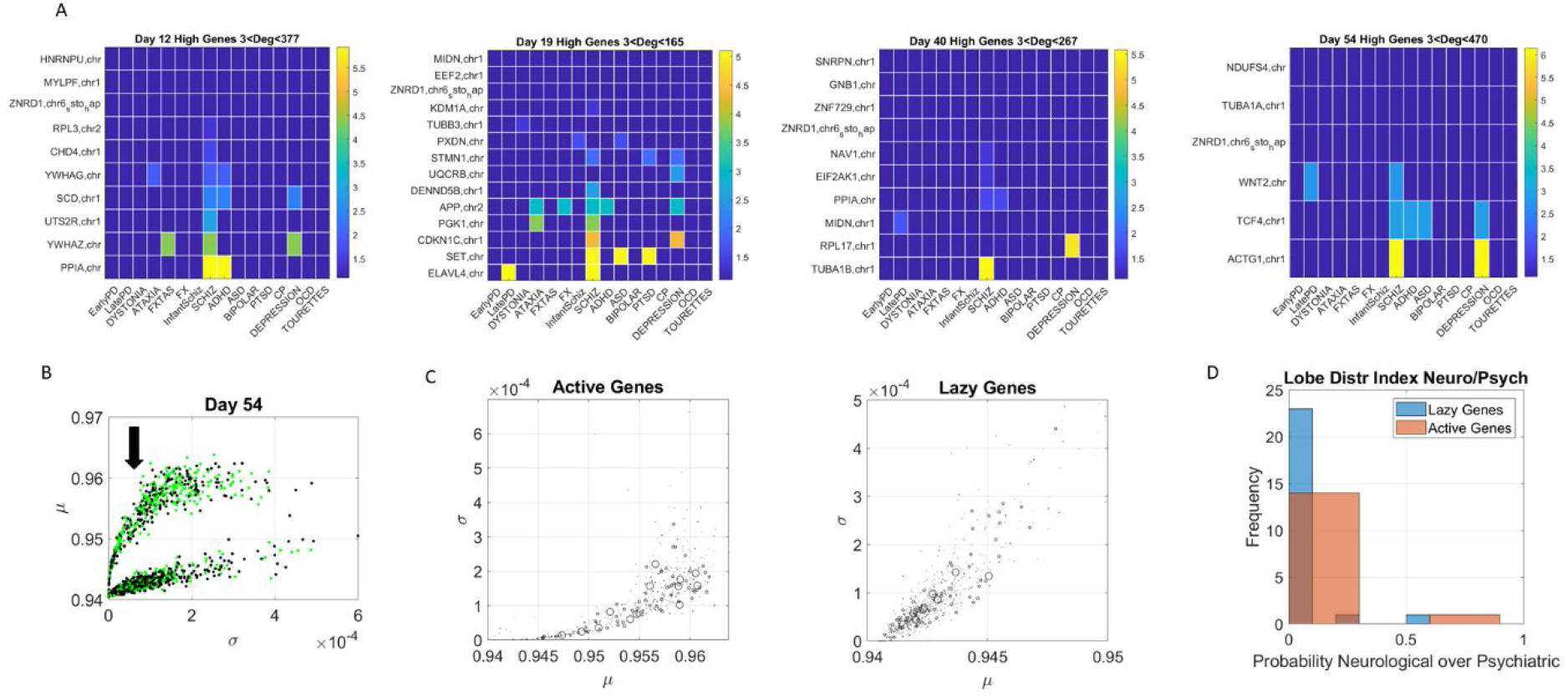
Evolution of genes’ degree distributions across neuropsychiatric and neurological disorders. (A) Days12-54 evolution of gene hubs (>=3degrees) on the curved lobe of active genes indicated in (B). Rows are genes and columns are disorders (neurological followed by DSM psychiatric.) (B) the arrow marks the lobe of active genes used to build the matrices. (C) Index *r* of equation 14 was obtained to ascertain the involvement of each disorder type per lobe. The marker size is proportional to this index, with larger markers representing more involvement in neurological disorders. Both lobes are represented and, despite the larger number of psychiatric disorders, a higher proportion of neurological involvement is featured in active genes. This is represented in the probability distribution graph in (D).

Similarly, for the lazy genes, we plotted the color map matrices involving disorders along the columns and genes along the rows. We see that in days 19 and 40 more genes partake as hubs and sub-hubs, with a more distributed network topology than in the case of active genes (as shown by the networks in Figures 10 and 11 for active and lazy genes, respectively.)

#### 6.3.1 Results from binary barcode (state space) and cells’ subtypes

Figure 15 shows representative results of tracking the gene’s state according to a binary code that sets the state ON if the node’s degree at a given day is above the average degree of the network that day, and OFF otherwise. With four readings we have 2^4^, 16 possible states, and each gene is assigned one possible state according to its network dynamics. In this figure (for simplicity) we plot two examples of genes with state [1111] remaining each day above the average degree of the network, taken across all nodes. Likewise, the example of [0100] is of a gene that only raises above the average degree of the network in day 19. The other configurations (dashed trajectories) on each plot reflect the trajectories of other genes in other binary state vectors. Using this information, we could build clusters of the cells that primarily had genes in a binary state. The use of both the fate and state dynamics of the gene enabled the tracking of the stochastic activity over time.

#### 6.3.2 Further similarities and differences between neurological and psychiatric disorders

Besides tracking genes expression and their variability in fate space along with the binary ON / OFF states of each gene and their involvement in neurological *vs*. psychiatric conditions, we selected the following clusters of tissues to evaluate genes’ involvement:

- Cortical Regions: Cortex, Frontal Cortex
- Neuromotor System: Caudate Basal Ganglia, Cerebellar Hemisphere, Cerebellum, Nucleus Accumbens, Putamen Basal Ganglia, Substantia Nigra
- Limbic System: Amygdala, Anterior Cingulate Cortex, Hippocampus, Hypothalamus
- Spinal Column: Spinal Cord Cervical C1, Nerve Tibial
- Glands: Adrenal and Pituitary gland

For each disorder in each clinical category (psychiatric or neurological) we removed the genes that would maximally express on tissues from the brain, the spinal cord, and glands, as described above. Upon removal we calculated (as in [9]) the resulting normalized changes in overall gene expression across these tissues. Then, we found in which percentile over all tissues each cluster of tissues belonged and identified the range of the change obtained within each cluster of genes (active *vs.* lazy). We computed the difference in percentile for each cluster of tissues to compare neurological *vs.* psychiatric disorders. We found statistically significant differences at the alpha 0.05 level (p<0.04, t-test.) Figure 16 shows the bar plots comparing the outcomes for active and lazy genes for psychiatric (red) and neurological (blue) disorders.

The psychiatric disorders included ADHD, ASD, Schizophrenia, Bipolar, Depression, Infantile Schizophrenia, PTSD and OCD. The neurological disorders included FXTAS, Early PD, Late PD, Ataxia, FX, Dystonia and CP and Tourette’s, given the involvement of the neuromotor systems in the last two disorders. Maximal differences in percentile change and in the reshuffling of disorders (which we plotted sorted by percentile in Figure 16) accounted as well for statistically significant differences between the tissues for the active *vs*. lazy genes (*p<0.04, t-test*).

## 7 Discussion

This paper uses genomic information to reframe a recently revived debate on the possible differentiation between neurological and psychiatric disorders. We reframed the question by addressing whether, despite a shared genomic pool, psychiatric and neurological disorders could be differentiated at very early embryonic stages. To that end, we developed and implemented a three-level approach that interrogated the human transcriptome trajectories of hundreds of hESCs as they reached neuronal state. Each level of inquiry offered new insights on the complex genetic origins of psychiatric and neurological disorders, highlighting fundamental differences between the two types of disorders. We uncovered and characterized two classes of genes with essentially different dynamics (distinct ON/OFF states) and fate (cumulative expression variability) featured throughout differentiation. Furthermore, using these classes of genes, we pointed at commonalities between the motor and limbic systems for one class (but not the other) that could possibly explain the current confounds in observational criteria. These analyses provide evidence supporting the notion that psychiatric disorders have substantial neurological underpinnings and yet their associated genes’ network interdependencies are significantly different at the early embryonic stages of cell differentiation. We next discuss each level separately.

### 7.1 Level 1: Marginal Distributions

Traditionally, transcriptome interrogation aims at uncovering different classes of cells with some genomic composition. This general approach tends to do away with genes that have low variability or asynchronous behavior, *i.e.*, they may be turned OFF in the initial stages, or have such low expression that their contribution is presumed negligible. We thought differently here and, instead of first trying to uncover cells’ classes, we transposed the cells x genes matrix and expressed each gene as a function of the cells’ expressions. Then the question was, for each gene, how was the gene’s expression across cells cumulatively contributing to the final neuronal fate. Furthermore, how was the gene’s state evolving across these different readings? We reasoned that some registered cells might not be the same from day to day, yet the expression of the genes would change across the transcriptome in some way that would lead to self-emergent patterns without discarding any genes. For each gene we obtained the marginal probability distribution of its expression to neuronal fate and measured across days the departure in expression variations, using the EMD metric appropriate to quantify differences or similarities between such frequency histograms.

Tracking this information without discarding any genes allowed us to visualize the genes associated with psychiatric and neurological disorders embedded in the full transcriptome. The new visualization tool revealed *two fundamental subtypes of genes* (as depicted in the two lobes of Figure 9.) Both lobes had a mixed composition of genes associated with psychiatric (according to the DSM-5) and neurological disorders. The question that emerged then was, what contribution was each set of genes (neurological *vs.* psychiatric) making to each lobe? We will defer that question to the third level of inquiry.

One lobe is characterized by genes of highly varying expression and high cumulative EMD whereas the other contains genes of low variance and low cumulative EMD. Here it may be worthwhile mentioning that traditional methods such as *PCA* and *t-SNE* would have likely missed the lazy genes of the second lobe, with lower expression and lower variability. And as we will see at the third level of inquiry, that would have missed an opportunity to capture the real evolution of these genes from the vantage point of probabilistic nodes interacting across a network.

### 7.2 Level 2: Co-dependent Genes

Moving on to a higher level of complexity, we studied the pairwise interactions between genes by focusing on the joint probability distributions of all possible pairs for each distinct group of genes associated with the various psychiatric and neurological disorders, according to sources found in the DisGeNet portal. By employing a robust statistical independency test, we were able to quantify the average degree of pairwise dependencies in each network of gene expressions. The initial degree of interdependence between genes associated with different pathologies (Figure 7A) allowed us to differentiate between neurological and psychiatric disorders.

Two main clusters appeared in this vector field, representing the increase in dependency from the initial to the final states. One cluster boasting a higher dependency increase was primarily neurological: early onset PD, Ataxia, Dystonia, FX and FXTAS. Yet, several DSM (psychiatric) disorders appeared at that level as well. These included disorders that are detectable in infancy, such as infantile schizophrenia, CP, OCD and Tourette’s. They share a highly compromised somatic-sensory-motor system and profound issues with the limbic system that without a doubt will impact the overall neurodevelopment of the individual. This pool of genes with high dependency gain across early embryonic stages of neuronal cell differentiation suggest rather early origins of such disorders and the highly interconnected evolution of their associated genes. They also showed a larger rate of change from the initial to the final state.

At a lower level of dependency increase, we saw mostly DSM psychiatric disorders, except for late onset PD. This is interesting considering the high incidence of dementia, hallucinations, and other altered mental states in late PD and other related tauopathies [27; 28]. These rates of change were more modest than those in the other group of disorders with a higher dependency increase. In particular, ASD, which lies approximately midway along this vector field, had the lowest dependency increase, signaling an altogether different signature of genes’ co-dependency during early embryonic stages of neuronal differentiation.

Indeed, ASD seems to lie on the border between the class of neurological and psychiatric disorders, which confirms at the genetic level that the spectrum of autism comprises pathologies of the nervous system that underlie its phenotypic conceptualization as a behavioral/mental disorder. This is supported by extended research showing biorhythmic patterns in autism with a unique signature of noise-to-signal ratio. This motor code is bound to impact the kinesthetic reafferent feedback from the periphery to centers of central control [10; 13; 29; 30; 31]. Here we observed the origins of such departures at these early embryonic stages of neuronal cells’ differentiation, whereby the genes associated to ASD manifested the smallest shift in dependency index value from the initial to the final state, accompanied by the smallest increase in genes’ statistical dependency. The change in dependency index (*i.e*., the range of values where the arrow denoting rate of change lands) overlapped with those ranges in FX, FXTAS, Ataxia, Dystonia and Early PD along the neurological disorders, and with infant Schizophrenia, along the psychiatric ones. This is interesting in view of recent work examining digital biomarkers of FX carriers who, despite their young age, manifested gait patterns present in Ataxia and FXTAS much like participants in the autism spectrum did [31; 32]. In this recent work, according to causal stochastic analyses, the kinesthetic feedback loops estimated from their gait largely departed (in both ASD and young FX carriers) from the neurotypical age-matched controls. They resembled instead the gait of patients with PD [32]. Our results here confirmed that the presence or absence of a disorder was not due to the mutation of a specific gene but rather resulted from the degree of co-expression among multiple genes. Therefore, in the context of the evolving human transcriptome, what truly separates neurological from psychiatric disorders is that the latter show much higher complexity of co-expression and interconnectedness in the early stages of cell differentiation, as compared to the genes associated with the former.

This hypothesis is reinforced if we observe the relative positions on this plane of Infant Schizophrenia and Schizophrenia, Schizophrenia and Bipolar Disorder, Tourette’s and OCD, PTSD and Depressions as well as ASD and ADHD, and ASD and Infant Schizophrenia. Once again, we have a transition in the initial degree of dependency from what characterizes Infant Schizophrenia as a neurodevelopmental disorder to what characterizes Schizophrenia as a disorder appearing in early adulthood. Also, OCD and Tourette’s have a high comorbidity, and symptoms of the latter may appear in the symptomatology of OCD. The same is true for Depression and PTSD, whereas both Schizophrenia and Bipolar Disorder belong in the psychotic spectrum of psychiatric diagnoses, according to the DSM-5. Hence, the proven clinical proximity in all three pairs recapitulates here in their proximity in our parameter space. Interestingly, ASD was once labeled as “Infantile Schizophrenia,” a clinical labeling that receives support in our parameter plane: ASD and Infantile Schizophrenia have nearly the same initial degree of dependency in our parameter space. Finally, Tourette’s is clustered among the psychiatric disorders. Indeed, there is ample debate on whether Tourette’s is a psychiatric or a neurological disorder, despite generally being classified as the latter [33; 34; 35].

When examining the cumulative changes in expression variability according to the EMD, a negative trend emerges in both types of disorders. The higher the genes’ variability accumulated toward Day 54, the lower the dependency index in this final state. However, ASD appears among the neurological disorders and Early and late PD appears among the psychiatric (DSM) ones (near to PTSD, Depression, OCD, CP, and Tourette’s). Furthermore, ADHD and Schizophrenia lie midway of this scatter and Infantile Schizophrenia and Bipolar disorders depart from the psychiatric group along the axis of variability, *i.e.,* their average cumulative EMD expressions are among the highest levels, along with those of the neurological disorders, at Day 54.

From these patterns, it may be possible to perceive that networks that reach a highly interconnected and complex state are characterized by genes that have more constancy in their statistical behavior. We revisit this proposition shortly, at the third level of inquiry. We reasoned here that, intuitively, this would make sense, since highly interconnected networks could dictate a gene’s behaviors in a distributed way whereas more disjointed networks would allow genes to behave in a more independent manner.

Our second level of inquiry considers pairs of genes, isolated from the rest of the network, and then integrates over all pairs to derive the degree of interdependency. We next explore a full probabilistic graphical model and visualize the global behavior of these genes. We characterize the topology of this network and identify the importance and role of different genes in the evolution of the network and subsequently on the origins of different disorders. To that end, we consider the multivariate probabilistic behavior of the two different subtypes of genes that we discovered at the first level of analysis.

### 7.3 Level 3: Genes’ Networks

By employing factorization techniques and graphical modelling, we were able to capture the evolutions of the networks of active genes (high variance, Figure 10) and lazy genes (low variance, Figure 11). In both graphical trees, we observed a hierarchical structure, with genes that were central (hubs) and associate with many other genes as well as genes that were leaf nodes. One key difference between active and lazy genes was that the latter have a less hierarchical organization on Days 19 and 40, with many small hubs and “clouds” of genes forming around the dominant hubs of the network. Note that the topology of the networks implies fragility, since removing a central hub or removing edges that connect large hubs would result in disconnected graphs, with the network of active genes being the most fragile of the two. We then catalogued the degrees of genes that were significant in each of the eight networks (two networks x 4 days for each genes’ class) and belonged to different neurological and psychiatric disorders (see Figures 12-14). Despite a vast majority of high degree genes being consistently associated with psychiatric disorders such as Schizophrenia (owing to the very high number of associated genes reported in DisGeNet), we found a far higher proportion of neurological involvement in active genes (Figure 12A and 12D. showing the probability distributions along with Appendix Figures 2–4). This result, along with those in Figures 13–14 for the lazy genes, pointed to a degree of overlapping of genes associated with neurological disorders with those associated with psychiatric disorders. This is captured as well on Table 1 of the Appendix, which we expanded in the Supplementary Material to catalogue the genes’ function, location, and phenotypes. The results also unambiguously separated genes involved in neurological disorders from those associated with psychiatric disorders in that the former were probabilistically more active (Figure 12D.) In contrast, the latter tended to be (probabilistically) lazier, yet forming more robust and distributed networks. From the network analyses and the index ratio quantifying neurological over psychiatric predominance, we concluded that, probabilistically, underlying psychiatric disorders are more lazy genes and underlying neurological disorders are more active genes.

**Figure 13.**
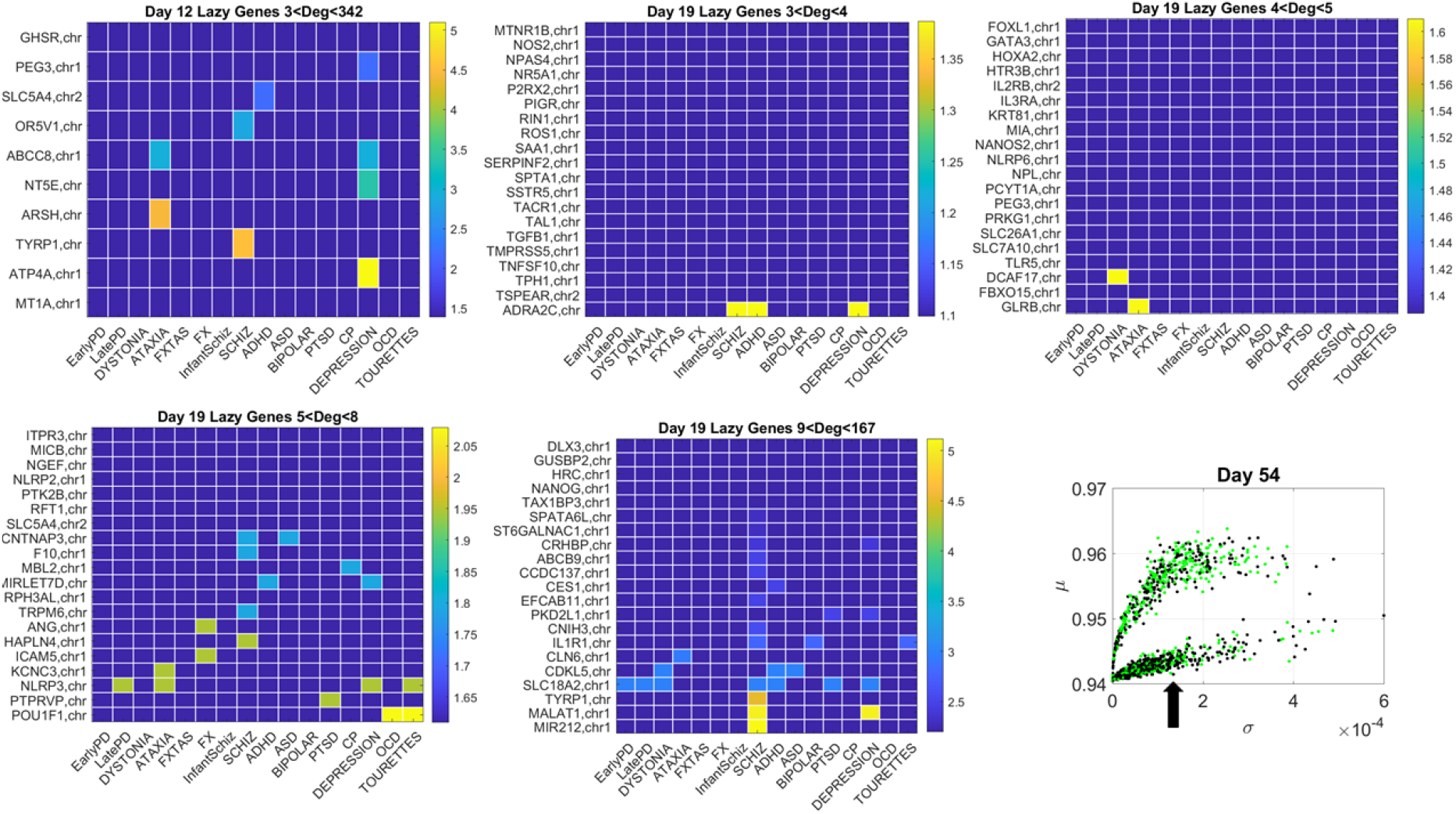
Tracking the lazy genes in the lower (line-like) lobe in Days 12 and 19 according to disorder (as in Figure 12) reveals the participation of several hubs in both neurological and psychiatric disorders. Arrow in the inset points to the lazy genes used to build these color map matrices and those in the next figure.

**Figure 14.**
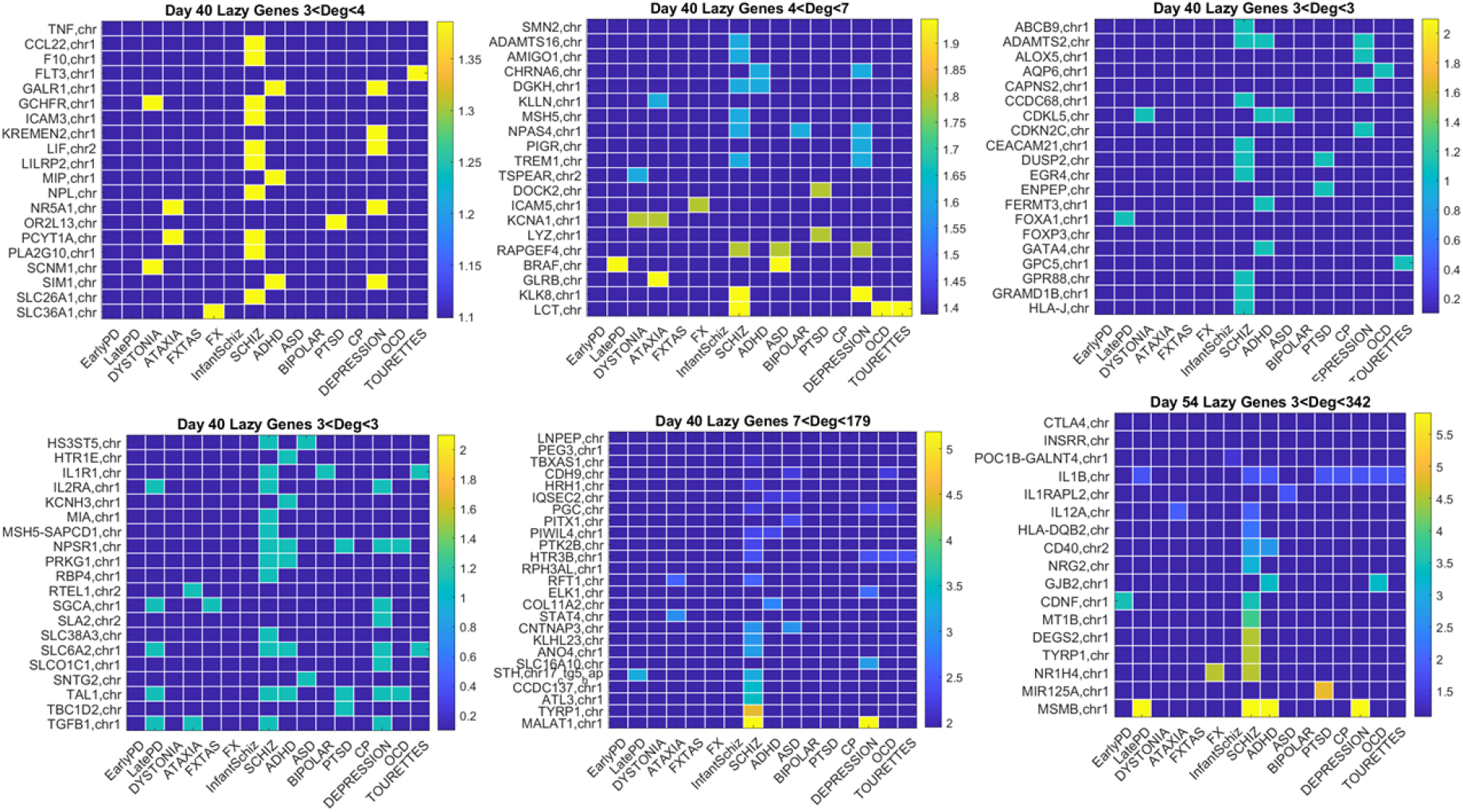
Additional lazy genes evolution in Day40 and Day54.

**Figure 15.**
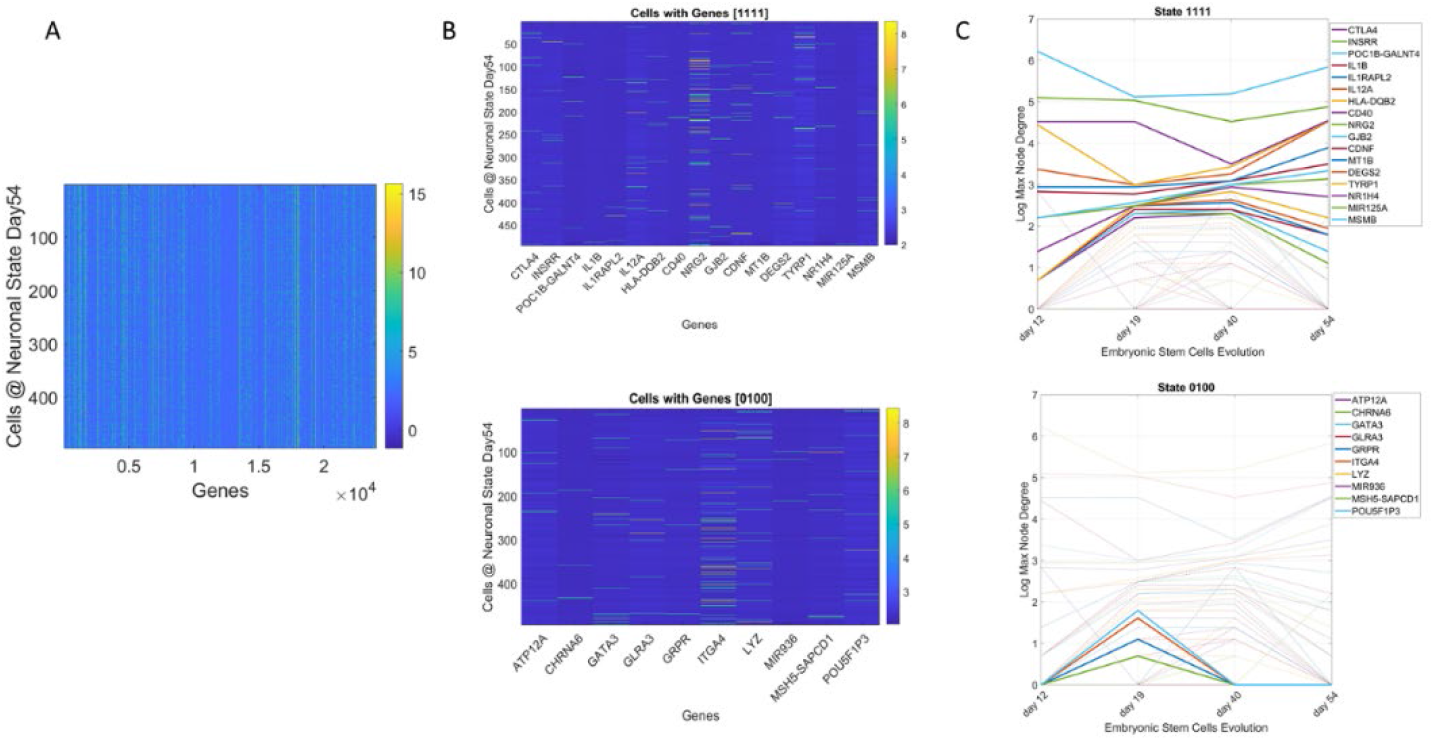
Genes’ states and cells’ hierarchical trees. (A) Matrix of 495 cells x 24,046 genes at day 54 of neuronal stages of hESCs. Each entry of the matrix is the counts in transcripts per million capturing the genes’ expressions at day 54. (B) Sample genes states [1111] and [0100] whereby the gene’s degree in the network is above the average degree each day (always ON, above connectivity threshold) and the case whereby the gene is OFF on day 12, then ON day 19 and OFF the remaining time. (C) Dynamic trajectory reflecting the log of the maximal degree of the node (gene) on the *y-axis* and time (days 12-54) under consideration on the *x-axis*. Dashed trajectories reflect the profile of all genes across binary space. Continuous lines on the top graph reflect the genes that remained ON above threshold (average degree) across all days *vs.* top graph genes with [0100] OFF/ON profile.

**Figure 16:**
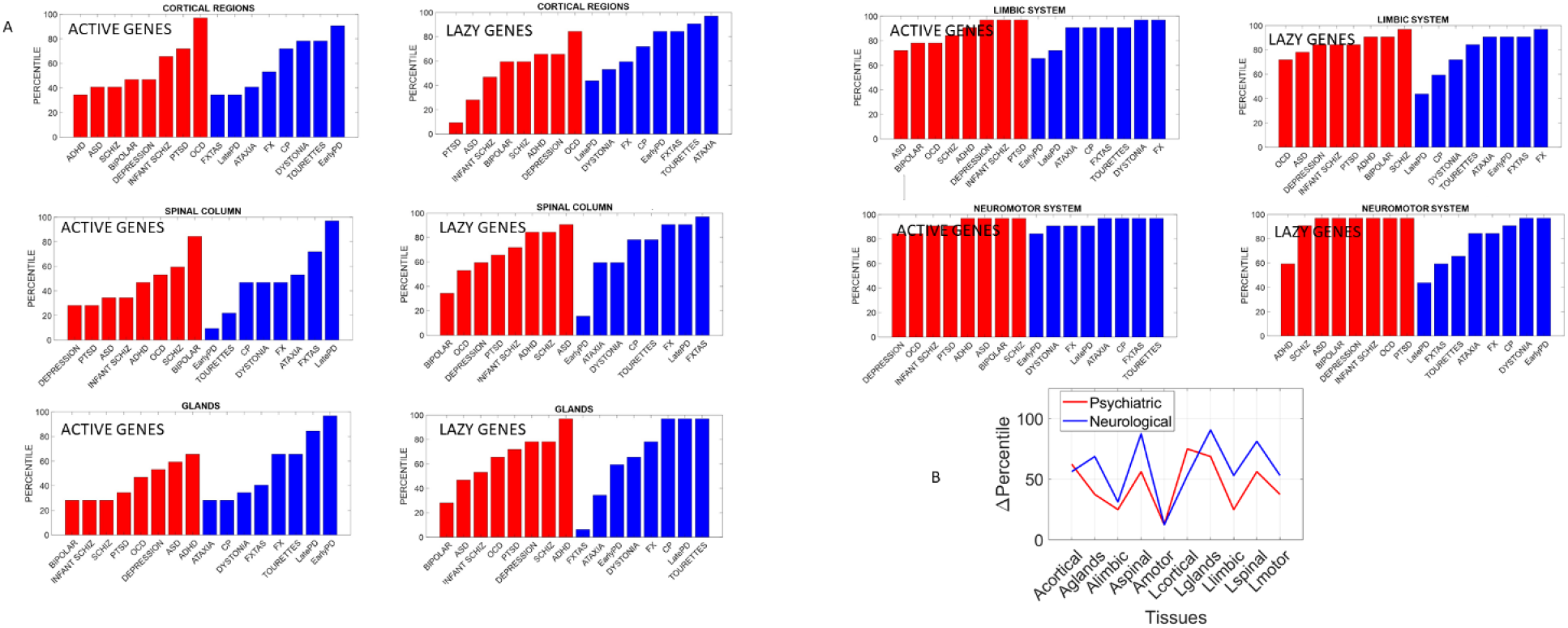
Differentiating psychiatric from neurological disorders by genes’ expression in tissues, for the active and lazy genes classes. (A) Percentile of change in genes expression on tissues for each disorder (red psychiatric *vs*. blue neurological) for tissues reported by the Consortium atlas of genetic regulatory effects across human tissues, GTEx v8 atlas of 54 tissues [26] here grouped into regions, cortical, spinal cord, limbic and neuromotor systems and glands (see text.) (B) Changes in percentile between psychiatric and neurological disorders are statistically significant at the 0.05 level and reflect the differences between active and lazy genes for each set of tissues comprising these regions of interest. The change was minimal in active genes for the limbic and motor tissues but appreciable in the lazy genes (often discarded.)

### 7.4 The Level of Average Genes’ Expression on Tissues

The fundamental differences quantified between psychiatric and neurological disorders using the different lobes of active *vs.* lazy genes extended to the tissues, as we probed these genes’ expressions on the 54 tissues from GTEx. Here we grouped tissues into brain regions (cortical, motor-subcortical and limbic), the spinal cord, and glands (see main text for details) and measured the percentile change between active and lazy genes upon removal of those genes in each of the disorders under consideration. We then grouped these disorders according to psychiatric and neurological clinical classification and compared the change in percentile between the two types of disorders.

Our analysis showed that, across different groups of tissues that typically serve different roles in the autonomic, central, and peripheral nervous system, fundamental differences emerged between the two types of disorders under consideration. The genes associated with psychiatric disorders expressed on these tissues differently than they did in neurological disorders. These differences were statistically significant between active and lazy genes in general. They were also statistically significant when considering the genes as part of psychiatric or neurological disorders. To that end we pooled the changes in percentile across all tissues and genes’ type of each disorder class (psychiatric or neurological) and found that the active *vs*. lazy genes classification served to separate psychiatric from neurological disorders. Furthermore, motor, and limbic regions were minimally different in active genes (relative to lazy genes) when comparing psychiatric to neurological disorders (Figure 16B.) This suggests that lazy genes, traditionally discarded, contribute to such statistically significant differences between psychiatric and neurological disorders. This result, paired with the probabilistically higher prevalence of neurologically associated genes across the active lobe and their presence in psychiatric disorders, supports the notion of motor involvement in mental pathologies. It is in this sense that one could argue that psychiatric disorders are also neurological disorders.

A main corollary of these results is that the lazy genes, which under traditional methods are likely discarded and excluded from the analyses, make an important contribution to the distinction between psychiatric and neurological disorders. This can be appreciated in Figure 16B whereby active genes (likely the ones entering the analyses under traditional techniques) do not separate motor and limbic systems between the two disorder types. It is the lazy genes that do so in the motor and limbic systems, and in other tissues as well. Further development of new analytical techniques that also include these genes with lower expression variability and asynchronous ON/OFF states may open new lines of inquiry across diseases in general.

### 7.5 Considering the Gene’s State Through a Binary Barcode

Interrogating the fate of the genes gave us a sense of ways to automatically cluster groups of genes according to the evolution of their expression variability. But, what about the changes in gene’s state? Once we reached the third level of inquiry and determined the gene’s degree of codependency at the network level, it was possible to examine the average degree of the gene to determine its hub activity level as above or below that average threshold (on a logarithmic scale to capture the non-uniform degree distribution.) In four days of measurements (in this case), we found genes that were always ON [1 1 1 1] in their hub networking role. These stood in contrast to those that were always OFF [0 0 0 0] in this role. Then, we found genes in other categories, thus building a barcode of binary states that allowed further examination of the genes embedded in the full transcriptome. Here we studied these pools of genes associated with neurological and psychiatric disorders, but the same type of method could be employed to interrogate the network’s state dynamics of genes associated with other disorders, with different frequency of recordings (here 2^4^ in four days of recordings, yielding 16 types that resulted in different cell classes expressing those main types.) However, other refinements of this method might offer new dynamics information with higher granularity of cell’s state that includes the low- and odd-varying genes with asynchronous states (such as the class of lazy genes uncovered here.) In this sense, further work is warranted to formalize a new embedding algorithm that considers the full transcriptome fate and state code to reveal topological invariants of the genes’ classes associated with known diseases.

### 7.6 Recapitulating Phenotypical Information Catalogued from Multiple Sources

We took our investigation one step further and compiled information from the OMIM online source for genes of degree > 3 of both gene lobes (active and lazy). Accordingly, we examined three categories of genes that, according to their association with both neurological and psychiatric disorders (overlapping roles), were associated with neurological disorders only and those associated with psychiatric disorders only. The “Neurological only” category consisted of only two such genes with degree higher than 3. Information about these classes of genes can be found in the Appendix Table 1 and in the expanded version of this table, including the inheritance of the identified genes and the functional properties of the proteins they encode in the Supplementary Material.

Both sets of genes associated only with psychiatric disorders and those associated with both the neurological and psychiatric ones were found to play critical roles in the developing nervous systems overall, fetal brain development, neurogenesis, neuronal differentiation, neuromodulation, synaptic organizations, healthy function of the senses, cortical development, and neuronal migration in the context of development. The psychiatric only-type genes were mainly associated with serotonin (5-HT), glutamate (NMDA) and nicotinic acetylcholine receptors (nAchRs), whereas the neurological and neuropsychiatric-type genes were associated with adrenergic (norepinephrine) and dopaminergic pathways. In both groups, specific genes were important for the GABAergic system.

In an interesting outcome (summarized in Appendix Figure 5), genes with a high network degree (large hubs) turned out to be crucial to the immune system. These genes are associated with autoimmune disorders and neurodegeneration, synthesis of growth factors, cancer and metastasis, inflammatory responses, and allergies. Notable cases are the active gene ELAVL4, with a network degree of 165. This hub is associated with Late Onset Parkinson’s disease and Schizophrenia and is related to paraneoplastic neurological disorders (PND) and autoimmune neuronal degeneration. Another gene, LIF, in the lazy-genes lobe, with a network degree of 4, has been hypothesized to play a functional role at the interface between the immune system and the nervous system. Here we recapitulated its association with Schizophrenia and Depression.

According to the OMIM literature, many of the identified high-degree genes (the hubs) play key roles in basic molecular and cellular functions, such as signal transduction, ATP synthesis, mitosis, cell-to-cell adhesion, intracellular signaling pathways, differentiation, proliferation, and transcription regulation. These large hubs also participate in these processes by influencing other genes’ functions (as predicted by the network’s topology) implicated in the formation of key components of the cell, such as the cytoskeleton and the extracellular matrix. These hub genes are crucial to the survival of the eucaryotic cell.

*These findings imply that the genes associated with neurological disorders are no more fundamental than the genes associated with psychiatric disorders or that are at the intersection of both disorders.* The origins of both types of disorders can be traced back to the embryonic stages of differentiation and development of the fundamental neural pathways and general biochemical cascades that characterize eucaryotic cell life. Moreover, in both types of disorders, key genes support the role and interaction of the immune system with the developing nervous system. The interaction between the two sub-systems is being explored and investigated by various researchers [36; 37; 38; 39], and their findings will shed light not only on the mechanisms of emergence of neurodevelopmental and neurological disorders but also on what the DSM-5 characterizes as “mental disorders.”

Amidst such heated debate on differentiating between neurological and psychiatric pathologies, perhaps, as synthesized by the OMIM literature behind their associated genes in our networks, a fundamental distinction is delineated by the neurotransmitter receptors and pathways linked to these genes. For example, the degeneration of dopaminergic pathways in the striatum is responsible for Parkinson’s disease-associated tremor, which is, at some point of the disorder’s progression, observable to the naked eye, thus shifting the focus to the “neurological” nature of PD (while ignoring the progressive dementia associated with it). In contrast, low serotonin in Depression leads to a “psychiatric” phenotype that sidelines motor control issues in these patients. On the other hand, high dopamine levels in Schizophrenia seem characteristic of this disorder, which despite being labeled “psychiatric” has a definite motor component [1; 14; 40; 41; 42; 43; 44].

### 7.7 Possible Utility of Reframing the Question of Psychiatric *vs.* Neurological Disorders from the Genomics Standpoint

Two important theoretical constructs could be proposed to further disentangle these disorders at the level of modeling system’s behaviors. One theoretical construct would have to borrow mechanisms from the immune/autoimmune systems to frame models of possible mechanisms. Given the synthesis of OMIM information, this line of inquiry will be important in future computational work.

Another avenue of theoretical inquiry explaining mechanisms to differentiate neurological from psychiatric disorders at the behavioral level would be related to the general framework comprising internal models of action (IMA) in the field of neuromotor control [45; 46]. We refer specifically to the “*principle of reafference*” [47] and related computational modeling. According to this basic principle, every time a movement is initiated by the nervous system, information is sent to the motor system and a copy of the signal is created, known as the *efference copy*. This enables the CNS to distinguish sensory signals stemming from (exogenous) external environmental factors from (endogenous) sensory signals coming from the body’s own actions [48]. According to the IMA, the efference copy is provided as input to a forward model, which predicts the sensory consequence of a motor command and measures the error between desired and attained outcome [45; 46]. Although IMA focuses only on error-correction and targeted-directed actions, complex movements are richly layered [49]. As such, endo-afference can be further <u>separated</u> into at least two components based on different classes of movements [50]. One type of endo-afference is classically associated with voluntary actions, *i.e.,* those deliberately aimed at a goal and operating under an error correction code [45; 46]. Another type of endo-afference, however, is associated with spontaneous or incidental actions, encompassing signals that transmit information about contextual variations associated with exploratory learning [51; 52]. Reafferent signals also include pain and temperature afference, a far more complex and elusive code that needs to be <u>distinguished</u> from the motor code in current biosignal’s analytics *e.g.,*as revealed in [53; 54].

The principle of reafference allows us to distinguish between different levels of *mental intent* [51; 55; 56; 57] and *physical volition* [14; 58; 59; 60]. This distinction can be conceptually mapped onto psychiatric and neurological issues, respectively, to inform hypothesis testing and theoretical modelling, possibly expanding criteria beyond subjective observation and opinion. However, we feel that the theoretical mechanisms stemming from immune/autoimmune systems will nontrivially add to our understanding of genomic differentiation between these classes of disorders. As such, they should be incorporated in a new internal model framework amenable to address different genes’ classes analogous to deliberate *vs.* spontaneous or error-corrective *vs.* exploratory modes of neuromotor control and learning, respectively [50; 52]. Here understanding recurrent loops of genes modulating other genes will be critical to forecast and track the onset timing of these disorders.

In summary, we have demonstrated at the level of hESCs that there are fundamental differences between psychiatric and neurological disorders when considering the full transcriptome. The inclusion in our analyses of genes associated with these disorders that nevertheless present odd or low variability and asynchronous ON/OFF states proved essential to making this differentiation. Considering only active genes would have missed this dichotomy. Furthermore, these distinctions extended to human tissues commonly studied in genomics. It is our hope that the multilayered, more inclusive approach offered in this work paves the way to open new lines of inquiry and help advance basic research in mental and physical health in general.

## 9 Authors contribution statement

TB analyzed data, designed analyses, wrote paper, SS curated data, edited paper, FHG supervised work, edited paper, TS supervised work, provided computational resources, EBT supervised work, conceived work, analyzed data, wrote paper. All authors contributed to the final state of the manuscript and agreed to its publication

## 10 Appendix

**Appendix Figure 1.**
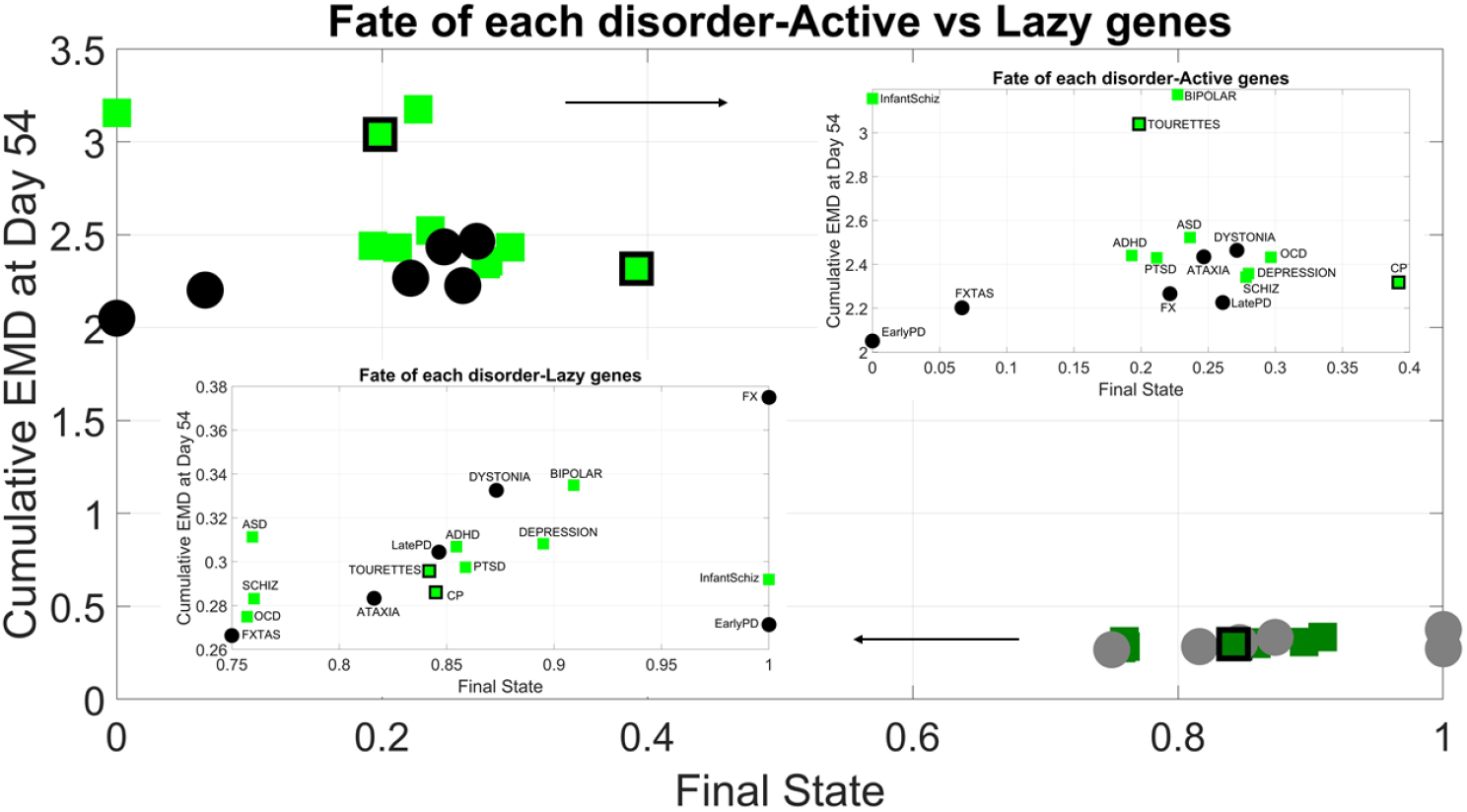
Unfolding disorders genes’ dependency index and fate by lazy vs active genes. Green squares represent psychiatric disorders. Gene squares with black edges represent psychiatric disorders that are also considered neurological. Active genes have higher cumulative EMD (fate metric) and lower dependency index at the final state. In contrast, lazy genes have lower expression variability according to the cumulative EMD and higher dependency index. Zooming in (insets) shows no discernable pattern among the disorders to differentiate in this parameter space the psychiatric from the neurological disorders.

**Appendix Figure 2.**
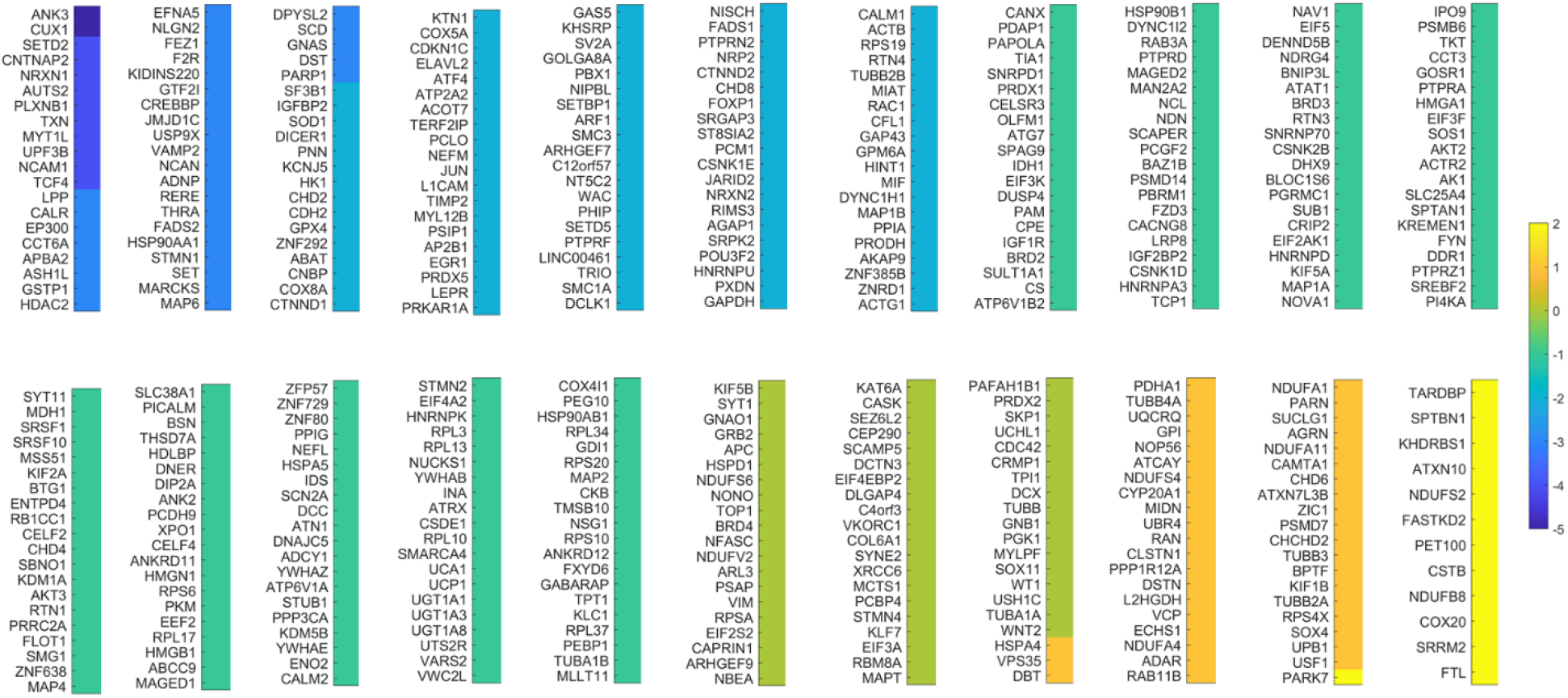
Colormap depicting genes sorted by disease index ratio (Equation 14) weighing the balance between active and lazy genes in neurological *vs*. psychiatric disorders. Active genes are plotted in the order of the scalar quantity (log of the ratio) whereby the higher the value (yellow), the higher the weight in neurological disorders.

**Appendix Figure 3.**
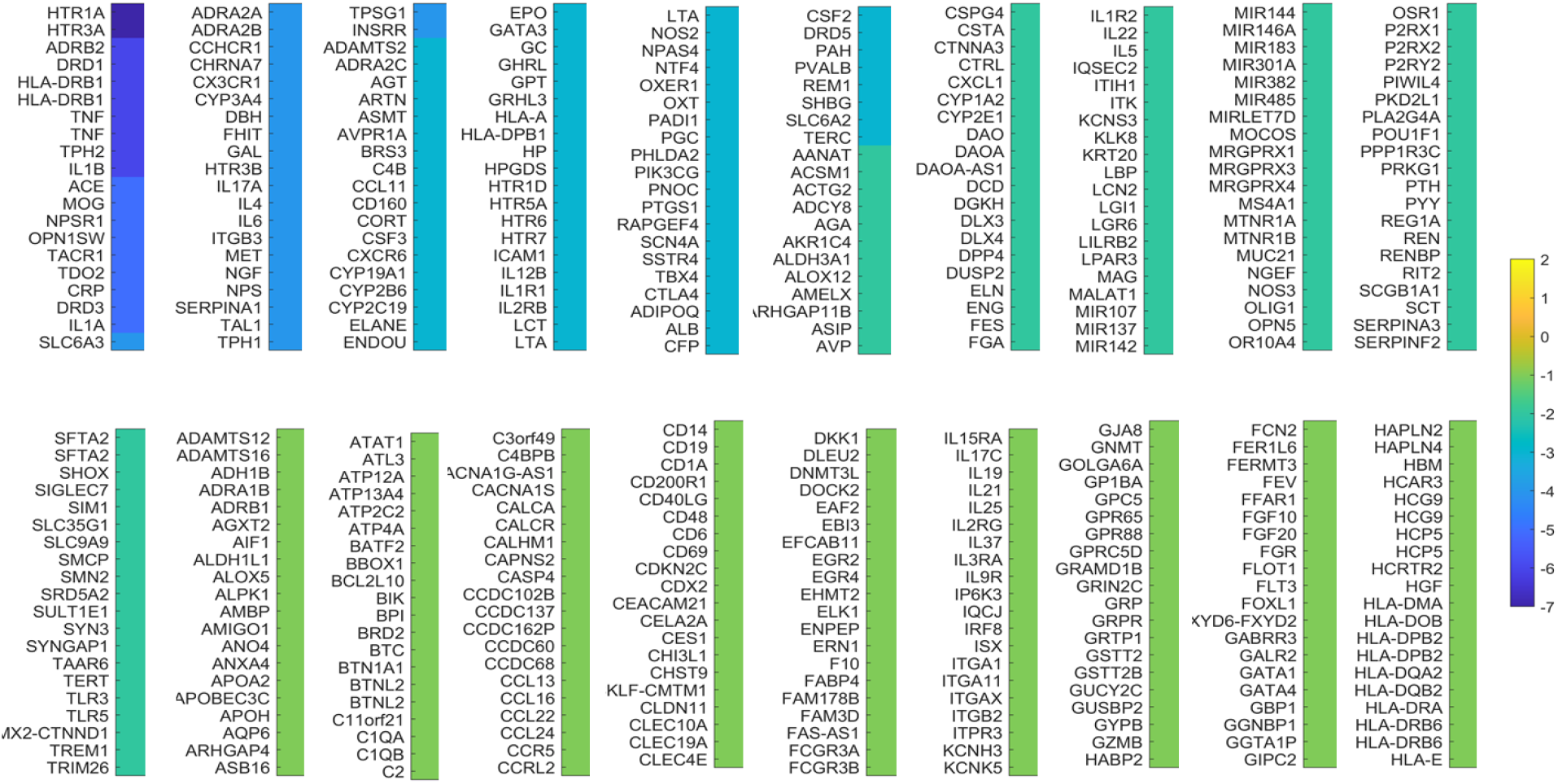
Lazy genes color coded as in Appendix Figure 1, based on the log of the disease index ratio.

**Appendix Figure 5.**
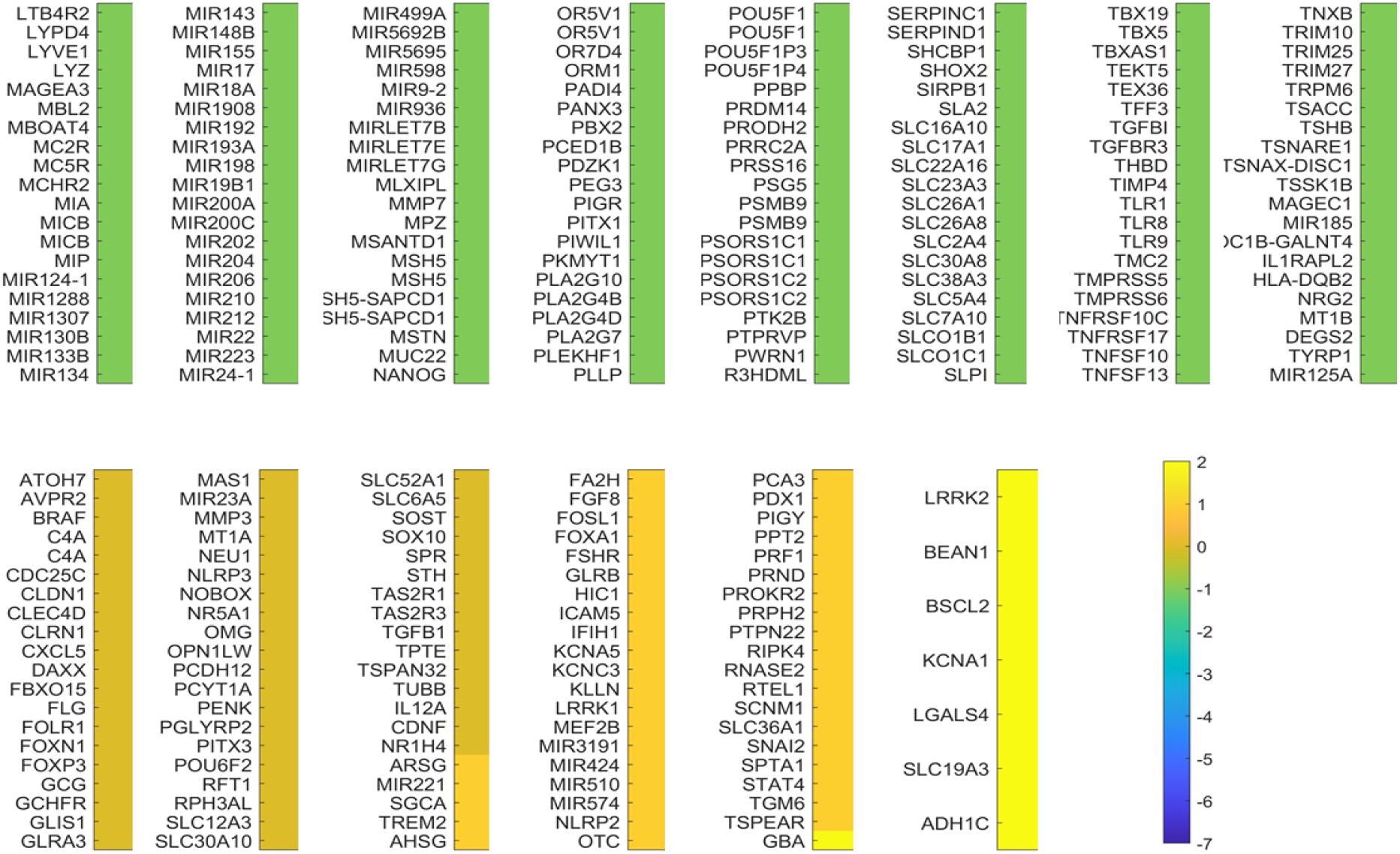
Lazy genes (continued) color coded as in Appendix Figure 1, based on the log of the disease index ratio.

**Appendix Figure 2.**
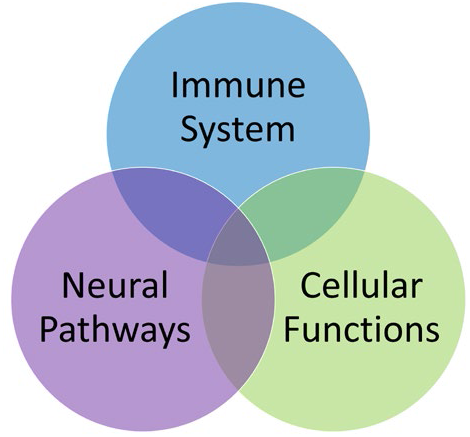
Venn diagram highlighting the three main clusters of genes identified by OMIM according to their function. A large number of genes play major roles at the interface between the immune system and the nervous system (See table in Supplementary Material). When considering psychiatric disorders alone or neurological disorders alone, very few high degree genes were found in neurological disorders alone. All three types of genes were found in genes common to both neurological **and** psychiatric disorders. For example, ELAVL4 of degree 165 was found in Late PD and Schizophrenia but it is also reported in paraneoplastic neurological disorders, and in autoimmune neuronal degeneration.

