## Supplemental Table for "Dynamic Interrogation of Stochastic Transcriptome Trajectories Using Disease Associated Genes Reveals Distinct Origins of Neurological and Neuropsychiatric Disorders"

| Neurological and Psychiatric | Genes | Degree | Day | Phenotype |
| --- | --- | --- | --- | --- |
| Ataxia, Schiz, ADHD  FXTAS, Schiz, Depression  Ataxia, Depression | YWHAG  YWHAZ  ABCC8 | 7  78  19 | 12 | YWHAG. Developmental and epileptic encephalopathy-56 (DEE56) is a neurodevelopmental disorder characterized by early-onset seizures in most patients, followed by impaired intellectual development, variable behavioral abnormalities, and sometimes additional neurologic features, such as ataxia (summary by Guella et al., 2017) Guella, I., McKenzie, M. B., Evans, D. M., Buerki, S. E., Toyota, E. B., Van Allen, M. I., Epilepsy Genomics Study, Suri, M., Elmslie, F., Deciphering Developmental Disorders Study, Simon, M. E. H., van Gassen, K. L. I., Heron, D., Keren, B., Nava, C., Connolly, M. B., Demos, M., Farrer, M. J. **De novo mutations in YWHAG cause early-onset epilepsy.** Am. J. Hum. Genet. 101: 300-310, 2017. [PubMed: [28777935](https://pubmed.ncbi.nlm.nih.gov/28777935/), [related citations](https://pubmed.ncbi.nlm.nih.gov/?cmd=link&linkname=pubmed_pubmed&from_uid=28777935)] [[Full Text](https://dx.doi.org/10.1016/j.ajhg.2017.07.004)]  YWHAZ  Popov et al. (2019) described a neurodevelopmental disorder characterized by global developmental delay apparent from infancy. Affected individuals had impaired intellectual development and poor or absent speech, as well as behavioral abnormalities. Most patients had significant facial dysmorphism, including coarse features, frontal bossing, and abnormal eye shape. Additional features were highly variable and included seizures, short stature, feeding difficulties, and skin abnormalities. For discussion of a possible association between this neurodevelopmental disorder and mutation in the YWHAZ Popov, I. K., Hiatt, S. M., whalen, S., Keren, B., Ruivenkamp, C., van Haeringen, A., Chen, M.-J., Cooper, G. M., Korf, B. R., Chang, C. **A YWHAZ variant associated with cardiofaciocutaneous syndrome activates the RAF-ERK pathway.** Front. Physiol. 10: 388, 2019. Note: Electronic Article. [PubMed: [31024343](https://pubmed.ncbi.nlm.nih.gov/31024343/), [images](https://www.ncbi.nlm.nih.gov/pmc/?term=31024343%5bPMID%5d&report=imagesdocsum), [related citations](https://pubmed.ncbi.nlm.nih.gov/?cmd=link&linkname=pubmed_pubmed&from_uid=31024343)] [[Full Text](https://dx.doi.org/10.3389/fphys.2019.00388)]  ABCC8 permanent neonatal diabetes neonatal diabetes, Babenko et al. (2006) screened the ABCC8 gene in 34 who did not have alterations in chromosome 6q or mutations in the KCNJ11 or GCK (138079) genes. In 2 patients with permanent neonatal diabetes, they identified heterozygosity for a mutation |
| Ataxia, FX, Schiz, ADHD, Depression  Ataxia, Schiz  Late PD,Schiz  Late PD, Ataxia, Depression, Tourette’s  Dystonia, ADHD, ASD  Early PD, Late PD, Dystonia, Schiz, ADHD, PTSD, Depression | APP  PGK1  ELAVL4  NLRP3  CDKL5  SLC18A2 | 20  45  165  7  20  20 | 19 | APP 21q21.3 Alzheimer disease 1, familial  PGK1 Xq21.1 Phosphoglycerate kinase 1 deficiency which catalyzes the reversible conversion of 1,3-diphosphoglycerate to 3-phosphoglycerate during glycolysis, generating one molecule of ATP  ELAVL4 The paraneoplastic neurologic disorders (PND) are a rare group of neurologic syndromes that arise when an immune response to systemic tumors expressing neuronal proteins ('onconeural antigens') develops into an autoimmune neuronal degeneration. PD  NLRP3 CINCA syndrome, also known as 'neonatal onset multisystem inflammatory disease,' or NOMID, is a rare congenital inflammatory disorder characterized by a triad of neonatal onset of cutaneous symptoms, chronic meningitis, and joint manifestations with recurrent fever and inflammation (Prieur and Griscelli)  CDKL5 deficiency disorder is characterized by seizures that begin in infancy, followed by significant delays in many aspects of development. Seizures in CDKL5 deficiency disorder usually begin within the first 3 months of life, and can appear as early as the first week after birth  SLC18A2 evidence that infantile parkinsonism-dystonia-2 (PKDYS2) is caused by homozygous mutation on chromosome 10q25 |
| Dystonia, ADHD, ASD  Late PD, Schiz, Depression  Late PD, FXTAS, Depression  Late PD, Schiz, ADHD, Depression, Tourette’s  Late PD, Schiz, ADHD, PTSD, OCD  Late PD, Ataxia, Schiz, Depression  Dystonia, Schiz  Ataxia, Depression  Ataxia, Schiz  Late PD, ASD  Ataxia, Schiz  Late PD, Schiz | CDKL5  IL2RA  SGCA  SLC6A2  TAL1  TGFB1  GCHFR  NR5A1  PCYT1A  BRAF  RFT1  STH Chr17ctg5 hap | 3  3  3  3  3  3  4  4  4  7  13  26 | 40 | IL2RA Immunodeficiency 41 with lymphoproliferation and autoimmunity, Diabetes, mellitus, insulin-dependent, susceptibility to  SGCA Muscular dystrophy, limb-girdle, autosomal recessive 3 17q21.33 autosomal recessive limb-girdle muscular dystrophy-3 (LGMDR3) is caused by homozygous or compound heterozygous mutation in the alpha-sarcoglycan gene (SGCA)  SLC6A2 Orthostatic intolerance 16q12.2 encodes a norepinephrine (noradrenaline) transporter, which is responsible for reuptake of norepinephrine into presynaptic nerve terminals and is a regulator of norepinephrine homeostasis (Kim et al., 2006) Kim, C.-H., Hahn, M. K., Joung, Y., Anderson, S. L., Steele, A. H., Mazei-Robinson, M. S., Gizer, I., Teicher, M. H., Cohen, B. M., Robertson, D., Waldman, I. D., Blakely, R. D., Kim, K.-S. **A polymorphism in the norepinephrine transporter gene alters promoter activity and is associated with attention-deficit hyperactivity disorder.** Proc. Nat. Acad. Sci. 103: 19164-19169, 2006. [PubMed: [17146058](https://pubmed.ncbi.nlm.nih.gov/17146058/), [images](https://www.ncbi.nlm.nih.gov/pmc/?term=17146058%5bPMID%5d&report=imagesdocsum), [related citations](https://pubmed.ncbi.nlm.nih.gov/?cmd=link&linkname=pubmed_pubmed&from_uid=17146058)] [[Full Text](https://dx.doi.org/10.1073/pnas.0510836103)]  TAL1 Autoimmune thyroid disease, susceptibility 8q24.22  TGFB1 19q13.2 Camurati-Engelmann disease, Inflammatory bowel disease, immunodeficiency, and encephalopathy Cystic fibrosis lung disease TGFB is a multifunctional peptide that controls proliferation, differentiation, and other functions in many cell types. TGFB acts synergistically with TGFA (190170) in inducing transformation. It also acts as a negative autocrine growth factor. Dysregulation of TGFB activation and signaling may result in apoptosis. Many cells synthesize TGFB and almost all of them have specific receptors for this peptide. TGFB1, TGFB2 (190220), and TGFB3 (190230) all function through the same receptor signaling systems  GCHFR 15q15.1 protein in rat that bound to GTP cyclohydrolase I (600225) and exhibited tetrahydrobiopterin-dependent inhibition of that enzyme. They found that this regulatory protein, which they termed GFRP, consists of a homodimer of 9.5-kD subunits and has a molecular mass of 20 kD. Milstien et al. (1996) used peptide sequences to clone the corresponding rat cDNA which encodes an 84-amino acid polypeptide. Northern blot analysis of rat tissues revealed that a 0.8-kb GFRP transcript was expressed at relatively high levels in liver and kidney and at somewhat lower levels in testis, heart, brain, and lung. Milstien et al. (1996) suggested that GFRP may play a role in regulating phenylalanine metabolism in the liver and in the production of biogenic amine neurotransmitters and nitric oxide  NR5A1 9q33.3 46, XX sex reversal 46XY sex reversal 3  PCYT1A 3q29 Spondylometaphyseal dysplasia-cone-rod dystrophy syndrome is characterised by the association of spondylometaphyseal dysplasia (marked by platyspondyly, shortening of the tubular bones and progressive metaphyseal irregularity and cupping), with postnatal growth retardation and progressive visual impairment due to cone  BRAF Noonan syndrome is a condition that some babies are born with. It causes changes in the face and chest, usually includes heart problems, and slightly raises a child's risk of blood cancer (leukemia). Noonan syndrome is a pretty common condition, affecting 1 in 1,000–2,500 babies. Delayed puberty  Down-slanting or wide-set eyes  Hearing loss (varies)  Low-set or abnormally shaped ears  Mild intellectual disability (only in about 25% of cases)  Sagging eyelids (ptosis)  Short stature  Small penis  Undescended testicles  Unusual chest shape (most often a sunken chest called pectus excavatum)  Webbed and short-appearing neck  RFT1 [3p21.1](https://www.omim.org/geneMap/3/372?start=-3&limit=10&highlight=372) Congenital disorder of glycosylation N-glycosylation of proteins follows a highly conserved pathway that begins with the synthesis of a Man(5)GlcNAc(2)-dolichylpyrophosphate (PP-Dol) intermediate on the cytoplasmic side of the endoplasmic reticulum (ER) membrane followed by the translocation of Man(5)GlcNAc (2)-PP-Dol to the luminal side of the ER membrane. RFT1 is the flippase enzyme that catalyzes this translocation (Helenius et al., 2002)  STH Chr17ctg5 hap Alzheimer Conrad, C., Vianna, C., Freeman, M., Davies, P. A polymorphic gene nested within an intron of the tau gene: implications for Alzheimer's disease. Proc. Nat. Acad. Sci. 99: 7751-7756, 2002. [PubMed: 12032355, images, related citations] [Full Text]  Verpillat, P., Ricard, S., Hannequin, D., Dubois, B., Bou, J., Camuzat, A., Pradier, L., Frebourg, T., Brice, A., Clerget-Darpoux, F., Deleuze, J.-F., Campion, D., the French Study Group on Alzheimer's Disease and Frontotemporal Dementia. Is the saitohin gene involved in neurodegenerative diseases? Ann. Neurol. 52: 829-832, 2002. [PubMed: 12447938, related citations] [Full Text] |
| Late PD, Schiz  Late PD, Schiz, ADHD, PTSD, Depression, OCD, Tourette’s  Ataxia, Schiz  Early PD, Schiz  Late PD, Schiz, ADHD, Depression | WNT2  IL1B  IL12A  CDNF  MSMB | 16  6  7  33  342 | 54 | WNT2 1p13.2 neonatal-onset chronic diarrhea O'Connell, A. E., Zhou, F., Shah, M. S., Murphy, Q., Rickner, H., Kelsen, J., Boyle, J., Doyle, J. J., Gangwani, B., Thiagarajah, J. R., Kamin, D. S., Goldsmith, J. D., Richmond, C., Breault, D. T., Agrawal, P. B. Neonatal-onset chronic diarrhea caused by homozygous nonsense WNT2B mutations. Am. J. Hum. Genet. 103: 131-137, 2018. [PubMed: 29909964, related citations] [Full Text] Ober, E. A., Verkade, H., Field, H. A., Stainier, D. Y. R. Mesodermal Wnt2b signalling positively regulates liver specification. Nature 442: 688-691, 2006. [PubMed: 16799568, related citations] [Full Text]  IL1B [2q14.1](https://www.omim.org/geneMap/2/552?start=-3&limit=10&highlight=552) Gastric cancer risk after H. pylori infection Interleukin-1, produced mainly by blood monocytes, mediates the panoply of host reactions collectively known as acute phase response. It is identical to endogenous pyrogen. The multiple biologic activities that define IL1 are properties of a 15- to 18-kD protein that is derived from a 30- to 35-kD precursor  IL12A 3q25.33 INTERLEUKIN 12A  CDNF 10p13 CONSERVED DOPAMINE NEUROTROPHIC FACTOR  MSMB 10q11.22 Prostate cancer, hereditary |
| Neurological Only |  |  |  |  |
| None |  |  | 12 |  |
| None |  |  | 19 |  |
| Dystonia, Ataxia | KCNA1 | 6 | 40 | KCNA1 12p13.32 Episodic ataxia/myokymia syndrome Potassium channels represent the most complex class of voltage-gated ion channels from both functional and structural standpoints. Present in all eukaryotic cells, their diverse functions include maintaining membrane potential, regulating cell volume, and modulating electrical excitability in neurons. The delayed rectifier function of potassium channels allows nerve cells to efficiently repolarize following an action potential |
| Late PD, FXTAS | SGCA | 3 | 54 | SGCA 17q21.33 Muscular dystrophy, limb-girdle, autosomal recessive 3 |
| Psychiatric Only |  |  |  |  |
| Schiz, ADHD, Depression | SCD | 11 | 12 |  |
| Infantile Schiz, ASD  Schiz, PTSD, Depression  Schiz, Depression  Schiz, Depression  Schiz, ASD, PTSD  Schiz, ADHD, Depression  ADHD, Depression  Schiz, ASD  OCD, Tourette’s  Schiz, Depression  PTSD, Depression  Schiz, Bipolar, Tourette’s  Schiz, Depression | PXDN  STMN1  DENND5B  CDKN1C  SET  ADRA2C  MIRLET7D  CNTNAP3  POU1F1  CRHBP  PKD2L  ILRL1  MALAT1 | 5  7  13  78  156  4  6  6  8  11  12  20  153 | 19 |  |
| Schiz, ADHD  Schiz, Depression  Schiz, ADHD, Depression  Schiz, PTSD  Schiz, ASD  Schiz, Bipolar, Tourette’s  Schiz, ADHD, PTSD, Depression, OCD  Schiz, ADHD  ADHD, Depression  Schiz, Depression  ADHD, Depression  ADHD, Depression  Schiz, ADHD  Schiz, Bipolar, Depression  Schiz, Depression  Schiz, ASD, Depression  Schiz, Depression  Schiz, OCD, Tourette’s  ASD, OCD  ADHD, ASD  Schiz, Depression, OCD  Schiz, ADHD  Schiz, Depression, OCD, Tourette’s  Schiz, ADHD  Schiz, Depression | PPIA  CRHBP  ADAMTS2  DUSP2  HS3ST5  IL1R1  NPSR1  PRKG1  GALR1  LIF  SIM1  CHRNA6  DGKH  NPAS4  TREM1  RAPGEF4  KLK8  LCT  CDH9  IQSEC2  PGC  PIWIL4  HTR3B  CNTNAP3  MALAT1 | 5  2  3  3  3  3  3  3  4  4  4  5  5  5  5  6  7  7  9  9  9  10  11  20  179 | 40 |  |
| Schiz, ADHD, ASD, Depression  Schiz, ADHD  ADHD, OCD | TCF4  CD40  GJB2 | 19  15  28 | 54 |  |
